# Machine Learning-Driven Optimization of Specific, Compact, and Efficient Base Editors via Single-Round Diversification

**DOI:** 10.1101/2025.07.25.666807

**Authors:** Mykyta Ielanskyi, Meng Wang, Lewis Scott, Lila Rieber, Stephanie Merrett, Johannes Schimunek, Andreas Mayr, Ian McDowell, Günter Klambauer, Tyson Bowen

## Abstract

Base editing shows great potential in research and clinical applications. Current iterations of the deaminases used to create precise single-nucleotide changes via base editing exhibit undesirable effects, including off-targeting, off-base editing, and bystander editing. Current deaminases are derived from either larger eukaryotic deaminases, which exhibit high levels of Cas-independent DNA targeting, or from evolved variants of the smaller E. coli TadA protein (ecTadA), which exhibits off-base editing. To overcome the limitations inherent to using a single protein sequence for engineering, we diversified newly identified TadA orthologs by DNA shuffling to yield millions of training sequences for measuring base editor efficiency. We trained generative models on the performance data from the pools of variants and drew on information-theoretic insights to efficiently explore the sequence space to generate diverse and high-performing deaminases. From a single round of diversification, we created a small set of novel and specific cytosine and adenosine deaminases that were markedly distinct in sequence from published base editor deaminases. We found that our model created deaminases generally outperform those we identified through typical directed evolution. The novel compact deaminases identified here show high on-base activity, comparable to the leading published base editors, and with demonstrably lower off-base activity.

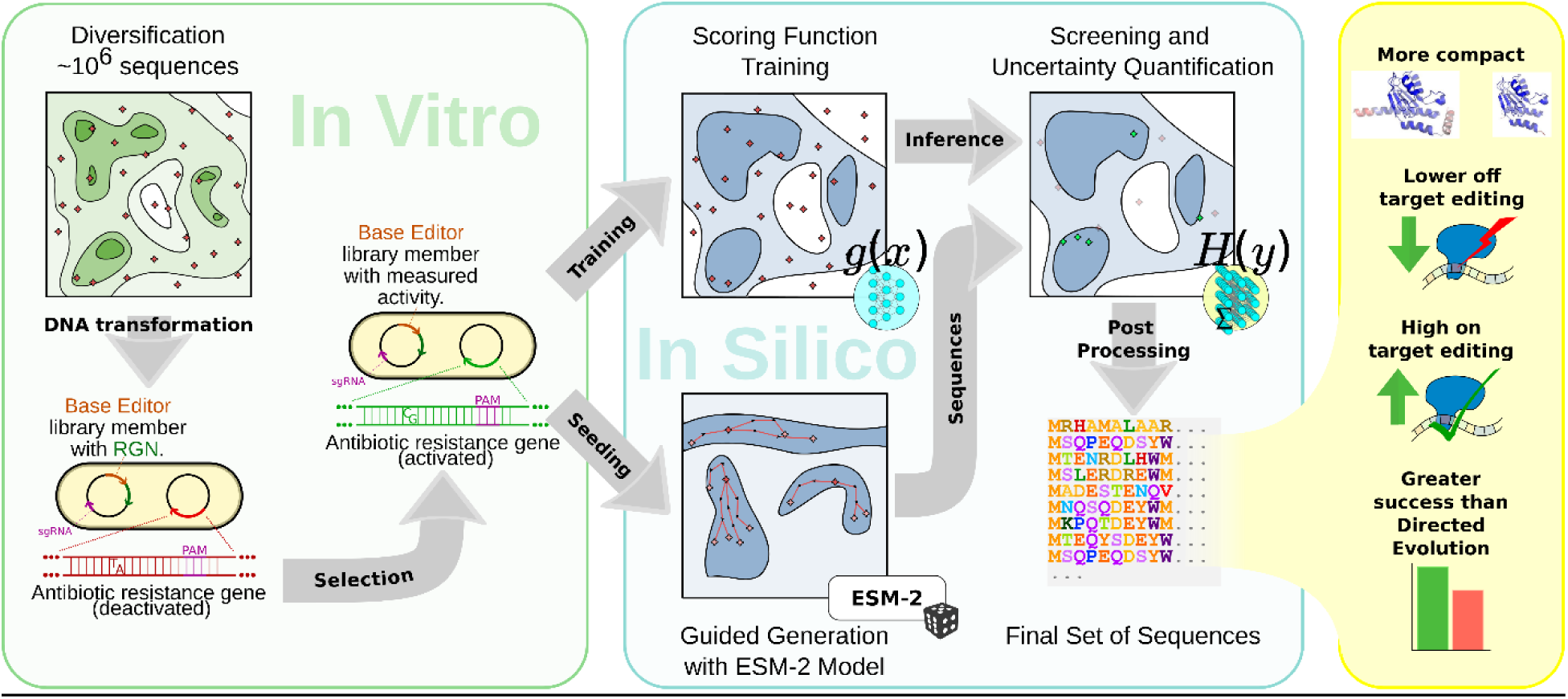
Graphical Abstract.

## 1 Introduction

Base editing is a gene-editing technology that combines the sequence-homing activity of a catalytically impaired CRISPR-Cas9 system with the enzymatic activity of a deaminase to enable precision editing free of potentially toxic double-stranded breaks (DSBs).(1–4) This new technology has the potential to cure diseases driven by point mutations, which are the largest known class of pathogenic mutations.(5,6) There are two major classes of base editors: cytosine base editors (CBEs), which catalyze C·G-to-T·A, and adenosine base editors (ABEs), which catalyze A·T-to-G·C.(3) CBEs typically feature naturally occurring cytidine deaminases, such as APOBEC, AID and CDA1, which natively act on single-stranded DNA.(7) To ensure minimal byproduct formation, uracil glycosylase inhibitor (UGI) is fused or simultaneously expressed to inhibit base excision repair and allow uracil to be replaced by thymine during DNA replication.(8) In contrast, ABEs feature evolved variants of adenosine deaminase acting on tRNA (ADATs), specifically TadA, from *Escherichia coli.*(*3,9–13*)

Despite high on-target DNA editing activity, base editors using natural or engineered deaminase variants still present problems such as Cas-independent off-target activity(13–16), bystander editing of non-target bases(17), and a large size relative to potential packaging vectors such as adeno-associated virus (AAV).(18) ABEs typically display many advantages for precision gene-editing over CBEs, including higher editing activities(11), lower Cas-independent off-target activities(9,11,13), and generally smaller sizes than CBEs.(19,20) Directed evolution (DE) studies have led to the development of improved adenosine deaminases that have expanded compatibility with CRISPR-Cas enzymes beyond SpCas9 and reduced bystander editing. Additionally, TadA has been evolved to function as a cytosine deaminase rather than an adenosine deaminase, as well as a dual adenosine-cytosine base editor (ACBE), as a workaround to the high rate of Cas-independent editing of naturally occurring cytosine deaminases. However, off-base activity, i.e., cytosine deamination by an ABE and vice versa, has been commonly reported in evolved TadA variants.(10,21) This drawback necessitates further optimization, especially for potential therapeutic applications. However, further optimization of TadA ignores the engineering potential within the natural variants of TadA orthologs, (22–24) which exhibit cytosine and adenosine base-editing activity with minimal engineering.(25)

Previous efforts to engineer ADATs to function as base editors have primarily relied on random or structure-inspired mutagenesis of ecTadA fused to SpCas9.(9–12,21,26,27) We hypothesized that this reliance on a single deaminase fused to a single Cas enzyme could be responsible for some of the known limitations of current TadA-evolved base editors, such as reduced activity when paired with non-SpCas9 RNA-guided nucleases (RGNs), deamination sequence context preferences,(11,26,27) and off-base editing.(1,23) Additionally, typical DE engineering approaches require multiple rounds of diversification and selection, where only a fraction of the total information from each round is considered and taken forward into the next.(28,29)

To address these limitations and overcome the information bottleneck between diversification rounds, we developed a single-step ML-based diversification approach to obtain a diverse set of highly active base editors. Generative deep learning approaches can identify biologically meaningful sequences by learning patterns from existing data, which enables the design of diverse, high-activity variants with fewer experimental iterations.(30–33) We started with a diverse set of newly discovered TadA orthologs, which we rationally modified to incorporate mutations homologous to those demonstrated to be effective in the engineered deaminases, ABE8e and TadA-CDd,(11,21) before performing DNA shuffling. Here, this strategy enabled the creation and testing of a larger sequence subspace than random mutagenesis while being populated by diverse regions of protein sequence with a high probability of retaining activity.(34), (35) These sequences were screened for deaminase activity, and the active samples were isolated and used to calculate enrichment values. This provided a rich source of data for training machine learning (ML) models to generate additional active deaminases through an *in silico* virtual screen. This allowed us to increase the rate of sequences that exhibited high eukaryotic activity to over 90%.

In contrast to other ML-guided protein engineering experiments (28,36), (37) which alternate between ML-guided diversification and DE cycles,(28) our approach uses a single step to diversify a set of proteins and predict new protein sequences that are likely to have high activity and fitness. To this end, our approach combined: i) a scoring function, that maps a sequence to an activity value and enables ranking, ii) a generative method that suggests biologically meaningful sequences, and iii) an uncertainty estimation method for selecting active sequences with a low false discovery rate and thus high precision. Our virtual screen yielded 28 diverse sequences with the highest predicted activity for both adenosine and cytosine deamination, which underwent further testing and characterization of base editing, specifically for on– and off-target activity. Our ML approach yielded a high number of distinct sequences with higher performance than those attained with DE alone, and improved specificity and comparable on-target activity relative to leading literature sequences. This new high-activity deaminase protein landscape brings high specificity and broad RGN compatibility, which can be explored to develop future performance gains for improved therapeutic strategies

## 2 Methods

### 2.1 Wet-lab processing

#### 2.1.1 Identification of Cas12f Systems

Novel Cas12f systems were identified as previously described.(38) Briefly, CRISPRCasTyper(39) was used to search public bacterial genome collections(40) for unknown Cas12f systems. The CRISPR operons of putative systems were cloned under control of the T7 promoter in the pET28 plasmid. The spacer sequence of three repeats of the CRISPR array was replaced with a specific target sequence. Synthesis was performed by GenScript. The PAM was identified in *E. coli* as previously described where active PAMs lead to plasmid cleavage and are depleted compared to controls.(38) Briefly, a plasmid library with the target sequence flanked by a random 8mer library was delivered to cells containing the expression plasmid and expression was induced. If the plasmid with the target sequence had a compatible PAM with the RGN, it was cleaved and the cells lost the antibiotic resistance marker on the plasmid. By comparing the relative frequency of different flanking sequences in the library pre and post depletion a PAM was determined.

Plasmid Interference assays were adapted from previous methods.(41) Reporter plasmids were synthesized on pTwist Amp High Copy or pTwist Chlor High Copy (Twist Biosciences) containing the same target sequences used in the PAM depletion assay with a defined PAM for each of the four Cas12f systems. The respective plasmids were then transformed into T7 Express E. coli (NEB) cells, and after a 1 hour recovery, the cells were plated in a 1:10 serial dilution onto LB agar containing IPTG, kanamycin, and the specific antibiotic for the targeting plasmid, either carbenicillin or chloramphenicol and grown overnight at 37C. The plates were compared to non-target control sequences that did not contain a matching spacer/PAM sequence.

Active Type V-F systems underwent bacterial RNA sequencing to identify their tracrRNA. In brief, the same expression plasmids used for PAM determination were expressed under identical conditions and RNA was prepared from *E. coli* as previously described.(38)

#### 2.1.2 Identification of ADAT Orthologs and sequence comparisons

A BLAST database was created from non-overlapping ORFs extracted from a public bacterial genome collection(42) and was used to perform searches on TadA sequences from *E. coli (ecTadA), S. aureus (saTadA), B. subtilis (bsTadA), S. typhimurium (stTadA), S. putrefaciens (spTadA), H. influenzae (hiTadA), C. crescentus (ccTadA),* and *G. sulfurreducens (gsTadA)* (Supplemental Table 11). Hits were extracted that were less than 70% homologous to ecTadA, or less than 85% homologous to other TadA sequences. Sequences were then filtered to only include those less than 220 amino acids long and that contain the prototypical deaminase fold and active site (D/H/C-[X]-E-[X15–45]-P-C-[X2]-C).(43)

The ADAT multiple sequence alignments from MAFFT FFT-NS-2(44) were used to create maximum likelihood phylogenetic trees in IQ-TREE.(45) Other sequence alignments were created by MAFFT E-INS-I(44) multiple sequence alignment in Geneious Prime (Biomatters Ltd) using ecTadA as a reference sequence. Predicted structures were generated using AlphaFold2(46) in Benchling, retrieved from https://www.benchling.com (2024). The structures are shaded by pLDDT values from 50 (red) to 100 (blue).

#### 2.1.3 Creation of Shuffled ADAT libraries

To create a starting pool of potential ABE deaminases, 95 candidate enzymes were identified (Supplementary Table 5) and aligned via a multiple sequence alignment to ABE8e and TadA.(11) Homologous residues for all introduced mutations from TadA to ABE8e were identified and mutated to the identified residue in ABE8e when possible (Supplementary Table 6). To create a starting pool of potential CBE deaminases, the same 95 candidate enzymes were identified and aligned via a multiple sequence alignment to TadA-CDd(21) and TadA. Homologous residues for all introduced mutations from TadA to TadA-CDd were identified and mutated to the identified residue in TadA-CDd when possible, as well as the V106W(11) mutation known to reduce Cas-independent off target activity (Supplementary Table 7). Sequences were then codon optimized for mammalian expression and synthesized by Twist Biosciences with a constant SV40 NLS sequence on the N-terminus and the constant XTEN32 linker on the C-terminus.(1)

These starting pools then individually underwent DNA shuffling(34) by being separately digested with eight different four-base-pair restriction enzymes (AluI, HaeIII, HhaI, HinfI, HpaII, MboI, RsaI, and TaqI) from New England Biolabs according to the manufacturer’s specifications (Supplementary Figure 6). The digests resulted in 3178 (ABE) or 2631 fragments (CBE), ignoring potential partial digestions. The digests were then pooled, and fragments from 75bp-300bp, representing approximately half of all the digestion products with no fragment being more than 65% of the size of the initial synthesis product, were isolated via gel electrophoresis and assembled via PCR into full-length deaminase constructs, which were isolated via gel electrophoresis and cloned into acceptor pET28 vectors synthesized by Twist Biosciences. For the ABE pool, the acceptor plasmids consisted of inactivated SpCas9(1) (dSpCas9_D10A_H840A), inactivated HoCas9(38) (dHoCas9_D10A_H566A), or inactivated enhanced LaCas12f (deLaCas12f_D93R_D240A_D416A). These three enzymes were chosen because they contained PAMs that could be utilized to target both the H193Y deactivating mutation of the Chloramphenicol resistance gene and the T89I deactivating mutation of the Spectinomycin resistance gene, and to select for deaminases that would work across a variety of RGNs.(3) The acceptor plasmids also contained targeting guides under T7 expression control to direct the RGN to the Chloramphenicol resistance gene, or multiple guides to target both the Chloramphenicol gene and the Spectinomycin resistance gene. Between the SV40 NLS and the XTEN32 linker on the acceptor plasmids was a unique ScaI recognition sequence. The plasmids were then linearized by digestion with ScaI-HF (New England Biolabs) before being assembled using NEB HiFi Assembly Mix (New England Biolabs) according to the manufacturer’s instructions. The assembled pool was then transformed into TOP10 cells (Thermo Fisher) to obtain approximately 5 million unique ORFs per RGN. For the CBE pool, the process was similar except the RGNs were additionally fused to BpSpUPP, a novel putative uracil-protecting protein identified from Bacillus phage vB_BpuM,(38) and contained targeting guides to direct the RGN to the H193R deactivating mutation of the Chloramphenicol resistance gene or the T89A deactivating mutation of the Spectinomycin resistance gene.(10) Approximately 5 million unique ORFs per RGN were again obtained.

#### 2.1.4 Bacterial Selection of Active Deaminases

Bacterial selection of active deaminases was adapted from previous methods.(3) Briefly, to identify functional deaminases, separate reporter plasmids were made for CBE or ABE activity. In general, pTwist Amp Medium Copy (Twist Biosciences) was synthesized to contain a deactivated chloramphenicol gene, or a deactivated chloramphenicol gene and a deactivated spectinomycin gene that could be reactivated by the proper deaminase activity. The mutations were H193Y (H193R) for ABE (CBE) activity in chloramphenicol resistance, and T89I (T89A) for ABE (CBE) activity in spectinomycin resistance. These reporter constructs were transformed into T7 Express *E. coli* cells (New England Biolabs) according to the manufacturer’s instructions. The cells were then grown to an OD600 of approximately 0.55 before being electroporated at 1800V with approximately 300 ng of purified library plasmid before being grown overnight at 37C to saturation in LB containing carbenicillin (100 ug/mL) and kanamycin (50 ug/mL). For the first round of selection, cells containing only the deactivated chloramphenicol gene were used with acceptor plasmids with a single guide RNA. Two independent selection transformations were performed for each RGN for the ABE screen and only a single transformation was performed for each RGN for the CBE screen. The following morning, the saturated growth was diluted 5% v/v into self-inducing Overnight Express™ Instant TB Medium (Millipore Sigma) containing carbenicillin and kanamycin to allow expression of the deaminase library and targeting RNA and grown for an additional 7h at 37C. The plasmids in the remaining starter culture were then extracted with a Miniprep kit (Qiagen). The entire culture was then plated onto agar plates containing carbenicillin, kanamycin, and chloramphenicol (25 ug/mL). A diluted plating onto non-selective LB carbenicillin and kanamycin was also done to obtain a count of the total colony forming units (CFUs) in the culture, as well as a diluted plating onto LB kanamycin with rifampicin (100 ug/mL) to obtain a count of the background mutation rate.(47) The plates were then incubated at 37C overnight, and the CFUs on each plate were counted to calculate the ratio of resistant CFUs to total CFUs. The colonies on the selective plate were then used to inoculate a fresh culture of selective LB media and grown overnight at 37C. The following day the plasmids were extracted with a Miniprep kit (Qiagen). The active deaminases were PCR-amplified, treated with DpnI (New England Biolabs) to remove residual plasmid DNA, inserted back into the empty RGN vector, and processed through the enrichment assay again. After the second round of enrichment, the selected deaminase library was moved into the acceptor plasmid with dual targeting gRNAs and underwent enrichment in cells containing the dual deactivated resistance gene plasmid. After the same growth and inoculation conditions, the cells were plated on selective plates containing either carbenicillin, kanamycin, and chloramphenicol, containing carbenicillin, kanamycin, and spectinomycin (100 ug/mL), or containing kanamycin, chloramphenicol, and spectinomycin. Surviving colonies were isolated from all growth conditions, and the resulting plasmids were isolated via miniprep (Qiagen). After the rounds, the initial deaminases pools and enriched deaminase pools were PCR-amplified and barcoded for next generation sequencing. Deep sequencing (2×300bp paired end reads) was performed on a NextSeq2000 (Illumina). Deaminase ORFs were extracted, counted, and normalized to total reads for each sample. Initially, paired-end reads were combined using FLASH2(48) to create unpaired reads. To minimize duplication caused by random insertions, deletions, and mutations related to library preparation and next generation sequencing, consensus sequences were clustered using MMSeqs2(49) in easy-cluster mode at 98% homology. The representative sequence from each cluster and cluster size, as the count of observations for each representative sequence, were used to assess enrichment by normalizing counts to total reads and calculating the log10 fold change of the final pool versus the initial pool. Enrichment was calculated as the log10 fold change of the final pool versus the initial pool. The top 20 unique enriched sequences for each RGN, as well as the top 20 overall enriched sequences were clustered at 85% (ABE) or 90% (CBE) homology to generate representative clusters. The top enriched sequence from each cluster was then chosen for advancement to eukaryotic testing as the directed evolution hit set.

#### 2.1.5 Dual Plasmid Lipofectamine Eukaryotic Activity Assay

The assay was performed as described previously.(38) Briefly, expression plasmids were synthesized for each selected base editing construct with dual NLS tags after codon optimization for human expression (Genscript) into the pTwist CMV plasmid (Twist Biosciences).(50) RuvC deactivated nickase versions of HoCas9, Nme2Cas9(51), and SpCas9 were used to test the deaminase constructs with a constant XTEN32 linker, and a C-terminal BpSpUPP was added to CBE editors. Targeted genomic sequences adjacent to the relevant PAM were selected and used to test the identified base editing construct (Supplemental Table 4). Respective targeting RNA was then synthesized under the U6 promoter on the pTwist CMV plasmid (Twist Biosciences) expressing a codon-optimized GFP. The plasmids were transiently transfected into HEK293T cells, and the eukaryotic activity assay was conducted following the method described in a previous study.(38)

Following transient transfection, the genomic DNA was extracted, and the relevant target region was amplified and prepared for next generation sequencing on a NextSeq (Illumina) following the manufacturer’s instructions for 2×150 reads. The reads were analyzed with CRISPResso2, and statistical comparisons were done in GraphPad Prism 9.2.0 (GraphPad Software, LLC) using a paired t-test when only considering a single target.(52)

When comparing across multiple targets, a linear mixed effects model(53) was used to assess the association of the deaminase and the on-base edit rate, with the target included as a random intercept. This approach allowed for estimating deaminase on-base editing while controlling for individual differences in target performance.

To determine the sequence context preference, we analyzed the frequency of each nucleotide located one position upstream of the on-base edit sites. Initially, we identified these edit sites within the quantification window of the CRISPResso2 alignments, which spans approximately 40 nucleotides centered around the guide. Next, we calculated the frequency of each nucleotide just upstream of the edit site by normalizing the total number of reads containing that nucleotide to the total number of reads mapping to the upstream site.

CRISPResso2 alignments were used to determine the frequency and position of on-base edits relative to the PAM. On-base edits were summed at each relative position and normalized to the total edits at that position.

#### 2.1.6 Cas-Independent DNA Off Target Editing

Orthogonal R-loop assays for Cas-independent off target editing were performed similar to established methods.(13) A catalytically dead version of *Staphylococcus aureus* Cas9 (dSaCas9) and corresponding SaCas9 sgRNAs that target genomic loci unrelated to the on-target site of the desired base editing were synthesized in identical plasmid backbones as the base editing constructs and guides. HEK293T cells were seeded in 24-well plates at 1.3 x 10^5^ cells per well in 500 μL growth medium (DMEM, 10% FBS, 1% Pen-Strep) and incubated overnight at 37°C, 5% CO_2_. In accordance with the manufacturer’s instructions, the cells were transfected with 200ng of plasmid containing targeting RNA (B2M_1), 200 ng of off-targeting SaCas9 gRNA,(13) 300 ng of plasmid containing dSaCas9 and 300ng of plasmid containing the effector protein using Lipofectamine 3000 (Invitrogen). The cells were then incubated at 37°C, 5% CO_2_ for 48 hours. Genomic DNA was extracted identically to the on-target assay, but both on-target and off-target regions were amplified using the Platinum SuperFi PCR Master Mix on the ProFlex 2 x 96-well PCR System before proceeding through the remaining barcoding and sequencing protocol and subsequent analysis with CRISPResso2. Statistical comparisons were performed as before.

#### 2.1.7 Cas-Independent RNA Off Target Editing

Cas-independent RNA off-target editing quantification was performed similar to established methods.(54) In brief, HEK293T cells were seeded in 6-well plates at 0.7 x 10^6^ cells per well in 2mL growth medium (DMEM, 10% FBS, 1% Pen-Strep) and incubated overnight at 37°C, 5% CO_2_. In accordance with the manufacturer’s instructions, the cells were transfected with 2500ng of plasmid containing targeting RNA (B2M_1), and 2500ng of plasmid containing the effector protein using Lipofectamine 3000 (Invitrogen). The plates were then incubated at 37°C, 5% CO_2_ for 48 hours. Cells were harvested and 1 x10^6^ cells from each sample were counted and sent to Azenta for RNA sequencing. Genomic DNA was extracted from each sample (1 x 10^5^ cells) for NGS analysis of the target site using the previously described protocol.

Fastq files were first trimmed using Trimmomatic v0.39 to “remove adaptor sequences, unpaired sequences, and low-quality bases”.(54) Next, reads were aligned to the hg38 reference genome with Hisat2 v2.2.1,(55) and the resulting alignments were sorted and indexed with SAMtools v1.13.(56) Variants were called with freebayes v1.3.6.(57) Indels were removed and the remaining point mutations were filtered to a minimum call quality of 20 using vcftools v0.1.16.(58) These filtered variants were summed across replicates with bcftools v1.13.(59) Any variants present in the null deaminase sample, representing the background rate of mutation, were removed from the base editor-treated samples.(15) Greater than 25 million reads were mapped for each sample.

### 2.2 Machine learning and computational modelling

Our ML approach contains three main components: a) a generative algorithm that suggests new proteins or mutations to proteins; b) scoring function that predicts activity/fitness based on a given protein sequence and c) uncertainty estimation algorithm that gives information about reliability of predictions of the scoring function. It builds upon the prior works on Reinforcement Learning-driven sequence design(60,61) and Bayesian Deep Learning.(62–64)

#### 2.2.1 Dataset and Preprocessing

##### Experimental Data

The CBE and ABE ADAT libraries that gave rise to the corresponding datasets were generated as described in Section 2.1.3. Because the single selection markers often saturated the assay, the counts from the two dual selection replicates were summed to obtain the training data. The statistics of the datasets for the ABE and CBE ADATs are given in Table 1. The reads per sequence were relatively low compared to several other DE assays from the literature,(37,65,66) but in general reads per sequence increased after selection, indicating that selection pressure was working. A fitness value corresponding to enrichment between the initial pool and after the final selection round was estimated for each unique sequence. Missing values were treated with pseudocounts (detailed in 2.2.2) or filtered away depending on the label preprocessing selected for the specific model.

**Table 1:** Summary of the dual selection data that was used to train the ML models.

| Base Editor | RGN | Pool | Reads | Unique Sequences | Reads per Sequence | Marker |
| --- | --- | --- | --- | --- | --- | --- |
| CBE | HoCas9 | R1 Pre | 67631 | 22687 | 2.981 | untreated |
|  |  | R1 Post | 1782672 | 440804 | 4.044 | ChlorSpec |
|  |  | R2 Pre | 714476 | 143965 | 4.963 | untreated |
|  |  | R2 Post | 2340647 | 431840 | 5.420 | ChlorSpec |
|  | LaCas12f | R1 Pre | 1116242 | 213156 | 5.237 | untreated |
|  |  | R1 Post | 1284387 | 370088 | 3.470 | ChlorSpec |
|  |  | R2 Pre | 475114 | 83773 | 5.671 | untreated |
|  |  | R2 Post | 2036743 | 287067 | 7.095 | ChlorSpec |
|  | SpCas9 | R1 Pre | 357623 | 114377 | 3.127 | untreated |
|  |  | R1 Post | 2537387 | 471502 | 5.381 | ChlorSpec |
|  |  | R2 Pre | 491957 | 160038 | 3.074 | untreated |
|  |  | R2 Post | 1640967 | 395878 | 4.145 | ChlorSpec |
| ABE | HoCas9 | R1 Pre | 27783 | 10206 | 2.722 | untreated |
|  |  | R1 Post | 2342696 | 644004 | 3.638 | ChlorSpec |
|  |  | R2 Pre | 854551 | 155015 | 5.513 | untreated |
|  |  | R2 Post | 1622237 | 341229 | 4.754 | ChlorSpec |
|  | LaCas12f | R1 Pre | 1518479 | 285553 | 5.318 | untreated |
|  |  | R1 Post | 2613712 | 375775 | 6.956 | ChlorSpec |
|  |  | R2 Pre | 2053521 | 392477 | 5.232 | untreated |
|  |  | R2 Post | 3221023 | 230801 | 13.956 | ChlorSpec |
|  | SpCas9 | R1 Pre | 180393 | 70023 | 2.576 | untreated |
|  |  | R1 Post | 1156268 | 397324 | 2.910 | ChlorSpec |
|  |  | R2 Pre | 584333 | 211590 | 2.762 | untreated |
|  |  | R2 Post | 919598 | 280631 | 3.277 | ChlorSpec |

#### 2.2.2 Pipeline Design

##### 2.2.2.1 Generative Model

To generate the protein sequence sets for ABE and CBE screening, we introduced an ESM mask filling approach. A seed sequence was first selected, and a prespecified number of amino acid residue positions were randomly and uniformly sampled across the sequence. For the sampled positions the residue tokens were replaced by mask tokens.

These masked positions defined the sites to be imputed, allowing us to generate plausible sequence variants through our iterative ESM mask filling strategy. To perform the imputation of the masks, we used the ESM-2(31) encoder, which has been trained in a BERT-style manner. At each iteration, ESM-2 produces a set of coupled per-position categorical distributions, each giving the predicted amino-acid distribution at a respective masked site conditioned on all observed (unmasked) residues and on the mask tokens occupying the remaining positions. Rather than choosing the next masked site at random, we concatenate all per-position logits into a single categorical distribution over position–residue pairs, imposing an artificial common scale that lets the model’s strongest preference determine the next update.

Concretely, let M denote the set of currently masked positions. For each I ∈ M, ESM-2 outputs a logit vector over its token vocabulary. We concatenate these vectors across all positions in M to obtain a single categorical distribution over position–residue pairs. Sampling from this distribution selects both the residue and the position to update; the corresponding mask is then replaced by the sampled token, and the partially completed sequence is re-evaluated with ESM-2. This procedure is repeated until all masked positions have been filled (see Supplemental Figure 18).

For each fill-in step, we retained the normalized categorical probability assigned to the selected position–residue pair. Aggregating these values yields a fill-trajectory pseudo-perplexity that we use as an uncertainty measure for the completed sequence.

While ESM-2 mask filling is designed to suggest point mutations, it is inherently unable to generate recombinant sequences. Since incorporating recombinant diversity is crucial for a broader exploration of the sequence space, we addressed this by using multiple seed sequences from the recombinant DE data. Specifically, we selected the top 32 best performing sequences for each of the three RGNs from the corresponding datasets to serve as seed sequences.

For sampling sequences, we specifically used the ESM-2(31) 650-million-parameter model. The number of injected mask tokens was between 1 and 3 to restrict sampling around the known clusters. We generated a pool of 10 million ABE and CBE candidate sequences for screening. Based on prior observations about protein language models, we expect this sampling procedure to bias against strongly protein-disruptive mutations.(67)

##### 2.2.2.2 Scoring Function

###### Sequence Encoding

We used one-hot encodings to represent each amino acid in the dataset because this performed better on the holdout set than using ESM-2 embeddings (data not shown). The ESM-2 embeddings would provide a more general representation capable of capturing information beyond the distribution of the sequences used to train the scoring functions, but might lose some domain-specific detail when compressing sequences into vector form. Relying on one-hot encodings seemed to have yielded better performance than relying on ESM embeddings, possibly because the ESM embeddings might have lost information relevant to the distribution of the ADAT data.

###### Use of Pseudocounts

The labels used for training were derived from enrichment factors (69), which in turn were computed from the original counts obtained from sequencing. Sequencing counts were processed using three pseudocount strategies: “no pseudocount”, “plus-one pseudocount” and “quantile plus-one pseudocount”. In the “no pseudocount” version, the sequences for which the enrichment score could not be computed directly were discarded. “Plus-one pseudocount” used a strategy of adding one to all counts. “Quantile plus-one pseudocount” added one to the 20% sequences with the lowest counts. To ensure comparability when evaluating model performance, the ‘plus￼one pseudocount’ and ‘quantile plus￼one pseudocount’ strategies used the same test and validation sets as the ‘no pseudocount’ strategy. We trained one scoring function per pseudocount processing strategy, each using its corresponding training set (which may differ due to discarding sequences). Because each pseudocount processing strategy may introduce its own bias, models from all three strategies were included in a combined prediction to average out strategy￼specific effects (for detailed model performances see Table 2).

**Table 2:**
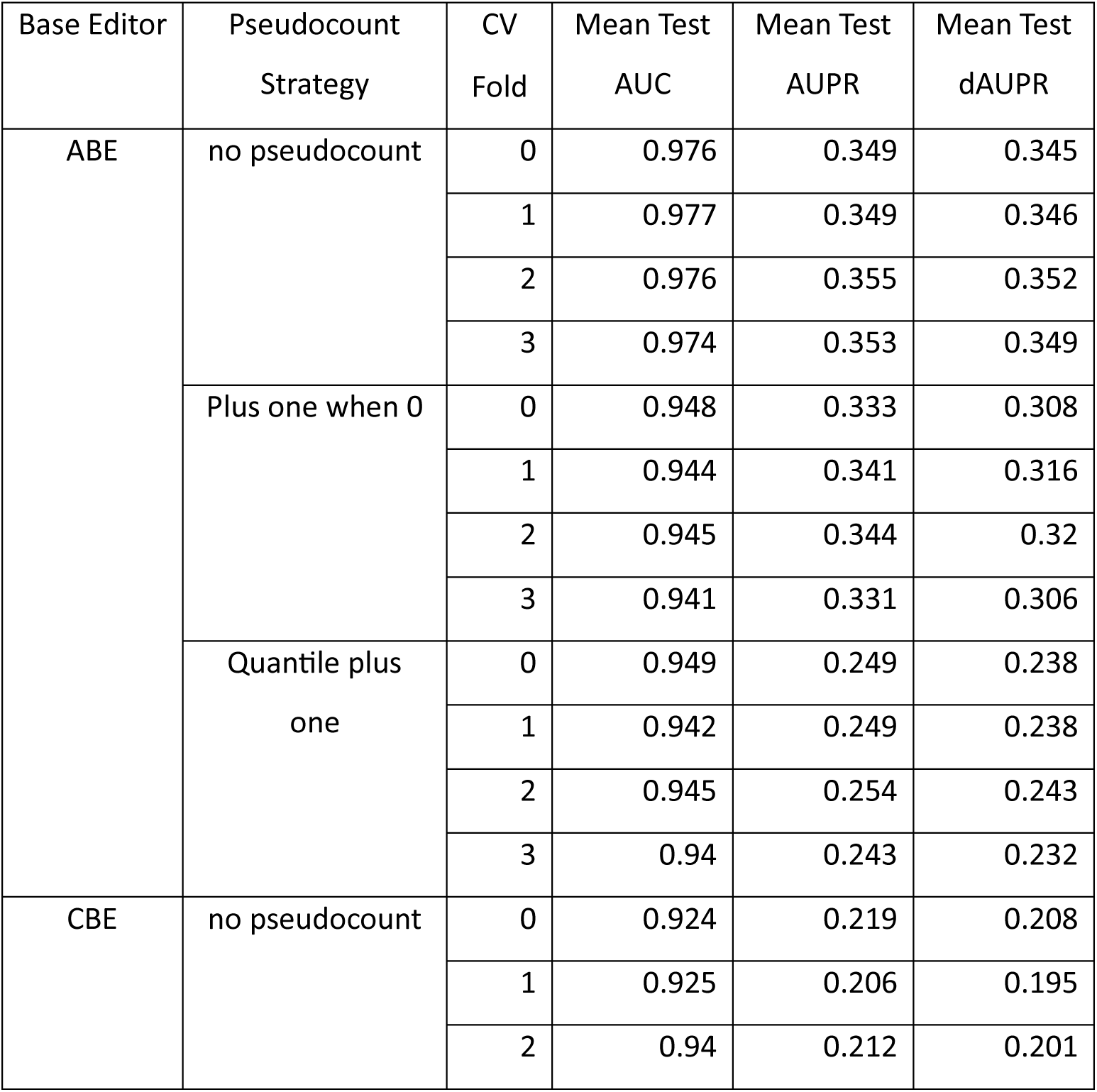
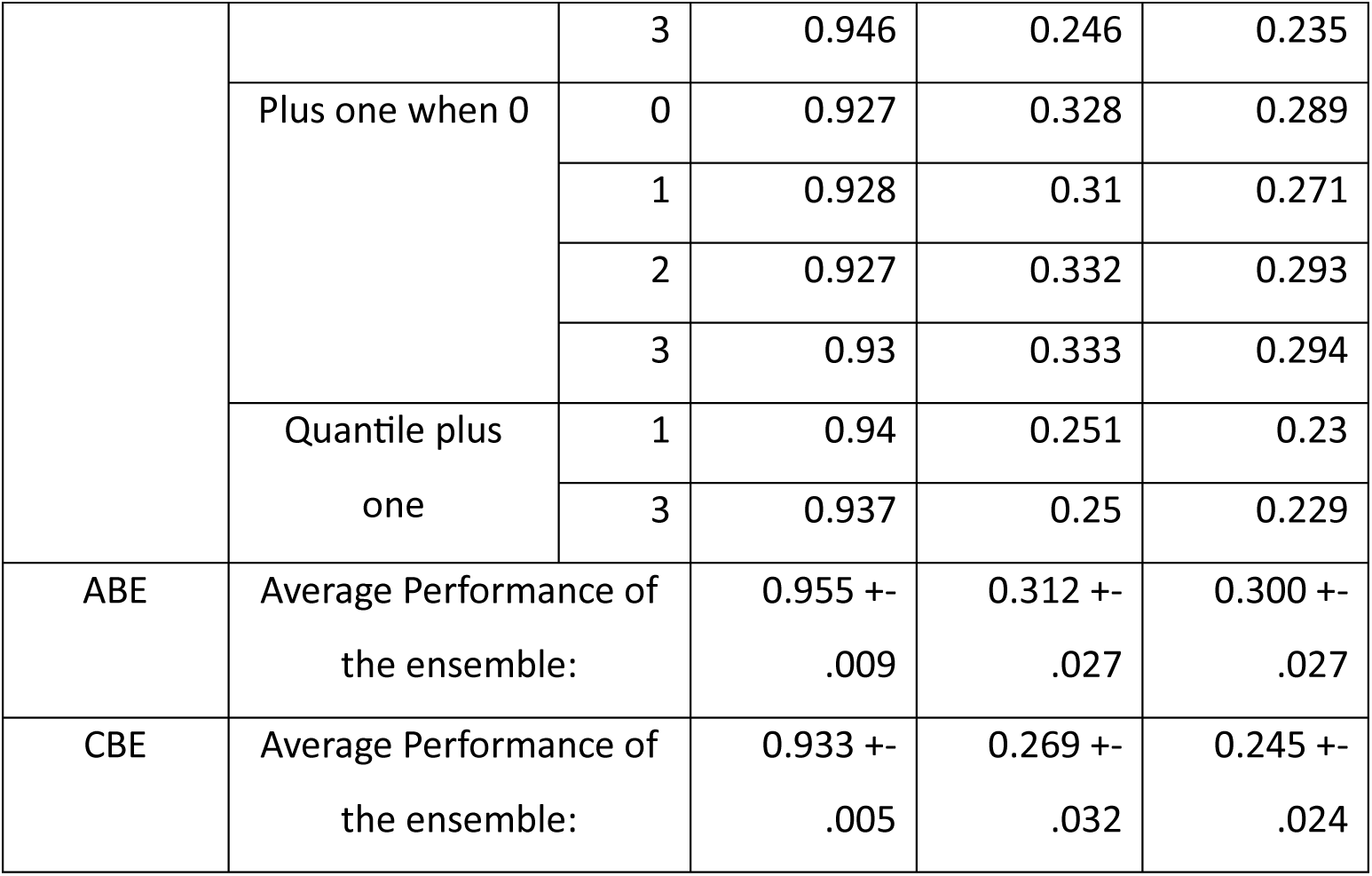
The performance of the ensemble of models used for scoring the sequences and uncertainty estimation, aggregated over all tasks and RGNs.

###### Training Algorithm

The change in relative abundance of a sequence between various conditions is measured by the enrichment factor. In the context of directed evolution, the enrichment can be computed between the rounds and represents an appealing target for modelling sequence performance in the absence of direct measurements of properties of interest. However, direct regression of this quantity is not straightforward, and requires additional normalization to address the high noise and imbalance present in the data. Several alternative methods exist such as using the asymptotic variance of the enrichment values derived from sequence counts,(68) or using a logistic regression approach to density ratio estimation for modeling the pre– and post-effect abundance of sequences in the pool.(69) Our ad-hoc training and evaluation scheme (Discretized Regression, DR) reformulated the regression task as a multitask classification problem by introducing several thresholds on the enrichment factor and converting the task into predicting, for each threshold, whether the value exceeds it. This threshold-based formulation, with one classifier per threshold, proved considerably more stable despite the imbalance (see Section 2.2.3).

For each threshold, we computed a class-weight factor from the positive-to-negative sample ratio and applied it to weight tasks accordingly during training. This helped moderate the effects of class imbalance specific to the classifiers. The following is the formulation of the DR objective:

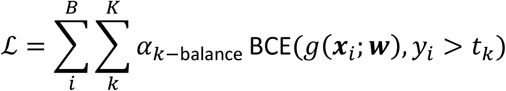

Here, we denote the predefined thresholds by *t_0_, …, t_K_*, ***w*** are the model parameters, **x**_i_ is the input sequence, y_i_ is an enrichment factor, and, B is the batch size. The class balance factor is denoted by *α_k-balance_* and BCE denotes binary cross entropy. The operation y*_i_* > *t_k_* provides binary labels indicating whether the enrichment factor y_i_ exceeds the threshold t_k_. Specifically, we have chosen thresholds of 1, 10, and 50. Our rationale here is that the lower-threshold tasks might provide signal relevant to learn meaningful intermediate representations, whereas the higher-threshold tasks should learn the signal to identify high-fitness sequences, which was subsequently used for ranking.

###### Model Selection

Model selection was performed using Cross Validation (CV). We used the same label definitions, which were used for training to perform model selection on the holdout sets. Enrichment thresholds 1x, 10x and 50x were ultimately selected for modelling by investigating the resulting class balance and batch size limitations. We employed standard metrics such as area-under-ROC-curve (AUC) and difference of area under precision recall curve (dAUPR), to assess the predictive performance across different architectures. The dAUPR on the holdout set was preferred as a selection metric due to the imbalance of the class labels and the priority we gave to retrieving positive examples correctly.(70) Our basic grid search hyperparameter tuning resulted in the selection of a three hidden layer 1D CNN with 128 channels in each layer with learning rate of 1e-2, weight decay of 1e-3 and Adam optimizer.(71) Adaptive max pooling was used after the 1D CNN stack to use the resulting representations in the linear output layer. The detailed performance of the final model ensemble can be found in Table 2.

##### 2.2.2.3 Uncertainty Estimation

Uncertainty estimation is important for determining whether a given input sequence is within the domain of applicability of the scoring functions trained on its training dataset. Epistemic uncertainty in formulation(72) is commonly used to this end:

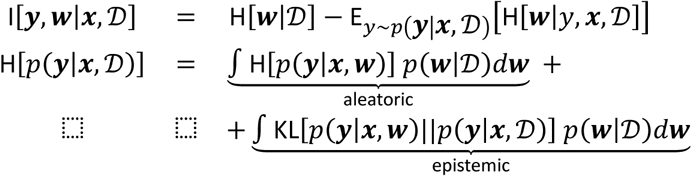

Here, p(y|x,D) is the Bayesian Model Average and p(w|D) is the model posterior. Houlsby et al. propose an approach that expresses information gain in terms of predictive entropies.(62)

Deep Ensembles is a popular and robust method that allows estimating uncertainty.(73) It uses a number of identically configured models trained from random initial weights to sample the posterior distribution. Ensembling deep neural networks has proven beneficial for predictive performance.(74)

The final scoring functions were ensembles of 1D CNN models (12 for ABE and 10 for CBE) that used one-hot encoding for data representation and DR for training. These ensembles included models trained with each of the three pseudocount preprocessing strategies across 4 folds. Models from all CV folds were combined to provide a bootstrapping effect. (31), (67)

#### 2.2.3. Scoring and ranking

We applied the model ensembles (Section 2.2.2.2 and Section 2.2.2.3) to rank all sequences generated by our generative models (Section 2.2.2.1). We selected the top 50,000 sequences for further analysis and processing. For these sequences we also calculated the epistemic uncertainty of the scoring function (see Section 2.2.2.3) and fill-trajectory pseudo-perplexity scores for the ESM generated sequences. The pseudo-perplexity should reflect the likeliness for a sequence generated by our iterative fill procedure, which is based on ESM-2’s token-level predictions with the intuition to measure how likely it is under the general distribution of protein sequences given the filling procedure. Large pseudo-perplexity values could indicate an unusual protein that might not be functional or able to fold. The epistemic uncertainty scores should point out sequences for which a reliable prediction cannot be made given the training dataset. For such sequences, the scoring functions might provide random values and have no predictive quality.(36,60) In other studies, the predictive quality of the scoring functions strongly decreased when there were more than three mutations relative to the training data.(68) To this end, we favored sequences with high predicted scores and low epistemic uncertainty.

For the final ranking, aggregate predictions for all ensembles were computed. The outputs of the heads trained to predict for t=50 (see Section 2.2.2.2) were used to rank the sequences. The predictions from the different RGNs were aggregated by multiplying their predicted probabilities. The top 200 unique sequences with the highest activity scores and lowest perplexity scores were selected from the SpCas9, HoCas9, and LaCas12f scoring results for both ABE and CBE. Each deaminase sequence was then aligned via a multiple sequence alignment to TadA and clustered. The computational mutagenesis did not produce any concentrated “hot spots” of novel mutations; rather, the introduced mutations were evenly distributed across the sequences. To reduce the number of potential sequences to test, representative sequences from each cluster were chosen that contained mutations in regions corresponding to residues 23-36, 46-51, 62, 73-83, 96, 106-127, or 142-166 of ecTadA but did not contain any mutations in the catalytic deaminase fold, corresponding to residues 57-59 or 86-90. The chosen regions were selected based on residues identified in previous studies(9–12,21,26) that increased activity upon mutation, but still included approximately 50% of the available sequence space. Mutations outside of these prioritized regions were allowed, but not used as a selection filter, and the selected mutations in the various clusters were distributed throughout the prioritized region (Supplemental Figure 8). The 28 highest-scoring sequences for ABE and CBE from these clusters were chosen for eukaryotic validation.

## 3 Results

### Identification and Characterization of Novel Cas12f systems

To identify highly active deaminases that can function flexibly across multiple RGNs, we designed our deaminase efficiency assay to test sequences across three RGNs and two different selection targets used for previous engineering efforts of TadA.(3) We opted to use two Type II systems (SpCas9 and HoCas9(38)) and a Type Vf system. Type Vf systems are compact (<700 amino acids), which enables packaging in AAV, and have been shown to function as base editors.(75) However, few have been reported to date, and those reported typically have lower activity than SpCas9, and none to our knowledge, with a compatible PAM that would enable targeting at the same selection targets as the Type II systems. Accordingly, we searched metagenomic sequences collected from diverse habitats(76) for undiscovered Cas12f enzymes using CRISPRCasTyper(39) and identified four novel Cas12f systems from the Clostridia class and are named each by their most identifiable phylogenetic group: LaCas12f (Lachnospiraceae), Ru2Cas12f (Ruminococcus), ChCas12f (Christensenellales), and CeuCas12f (*Coprococcus eutactus*). LaCas12f, has a 5’ TC PAM (Supplemental Figure 1) that can target the same mutations as the Type II systems used for our engineering effort (Supplemental Figures 2 and Supplemental Tables 1-2). To confirm robust bacterial activity necessary for our engineering assay, we performed a plasmid interference assay to demonstrate high activity of the systems.(41) We observed successful depletion with all four systems in *E. coli* (Supplemental Figure 3). We then performed small RNAseq to identify the tracrRNA. The identified Cas12f systems all possessed and identifiable tracrRNA, which was immediately followed by an antirepeat region immediately upstream of the CRISPR array (Supplemental Figure 4). The entire transcript region from the start of the expression profile through the spacer was dubbed a long monomeric nucleic acid targeting RNA (lmntRNA). We then sought to characterize the eukaryotic editing potential of these systems, but observed poor performance in HEK293T (Supplementary Tables 3-4). The activity of the novel systems was marginally improved by identifying residues identical to those previously introduced via multiple sequence alignments to create an enhanced version of Un1Cas12f1 (CasMINI; D143R_T147R_E151A).(75) D143 was found to be conserved in all newly identified Cas12f systems and mutated to arginine in all contexts to create LaCas12f_D93R, Ru2Cas12f_D82R, ChCas12f_D91R, and CeuCas12f_D82R. The lmntRNA was improved to create an enhanced lmntRNA (elmntRNA) by truncating down the antirepeat:repeat stem loop and introducing a GAAA tetraloop(75), both approaches improved activity, however overall eukaryotic editing activity remained low (Supplementary Tables 3-4). However, the robust activity in *E. coli* allowed us to proceed with our engineering strategy.

### Bacterial Selection of Active Shuffled Deaminases

The same metagenomic database used to identify the Cas12f systems was also searched via BLAST with eight different known TadA orthologs to identify a diverse starting pool of 95 likely deaminase sequences for DNA shuffling (Supplemental Figure 5 and Supplemental Table 5). For the search we selected the original TadA sequences from *E. coli (ecTadA)* used to make an adenosine base editor ,(9) as well as the ortholog from *S. aureus (saTadA)*,(77) which has also been extensively studied. The remaining six TadA orthologs were chosen to be diverse in sequence space, with between 40-61% identity to each other, while still possessing sufficient regions of homology to allow for successful DNA shuffling, and to be similar in length. Homologous residues to those identified in the creation of ABE8e or TadA-CDd were identified and mutated to the selected residue in ABE8e or TadA-CDd when possible, to create a starting population for ABE or CBE DNA shuffling (Supplemental Table 6&7).(11,21) In DNA shuffling, the starting sequences are digested into small fragments, which can then recombine via small sections of microhomology to generate a large pool of diverse sequences with an increased likelihood of maintaining desired characteristics and function.(34,35) The range of homology between the newly identified sequences allowed for multiple opportunities for successful microhomology crossover at multiple independent sites of the primary sequence.

We then incorporated our pool of millions of shuffled TadA orthologs into the three RGNs and tested their performance in a selective bacterial enrichment assay in which on-target base-editing activity rescued an antibiotic resistance gene (Figure 1a). By comparing the number of resistant CFUs formed under the selective deaminase conditions to those formed under non-selective or non-targeted conditions, we were able to quantify the effectiveness of our pool after each successive enrichment. For the first round of enrichment, there were elevated levels of targeted active deamination compared to a dSpCas9 with no deaminase (Figure 1a). It should be noted that the inclusion of BpSpUPP may have increased the background mutation rate of the CBE pool, as inhibiting uracil glycosylase is known to be mutagenic.(78) Because the second round of enrichment saturated our detection limit across most of the conditions, an additional antibiotic was included for the final round of enrichment. This final round of enrichment saw saturating activity on both single antibiotics in most conditions as well as substantial activity in the dual selection condition, indicating that we had enriched for a highly active pool of deaminases. Additionally, the off-target mutation rate increased noticeably over the dSpCas9 control in the third enrichment round, indicating a potential increase in non-targeted active deamination activity.

**Figure 1.**
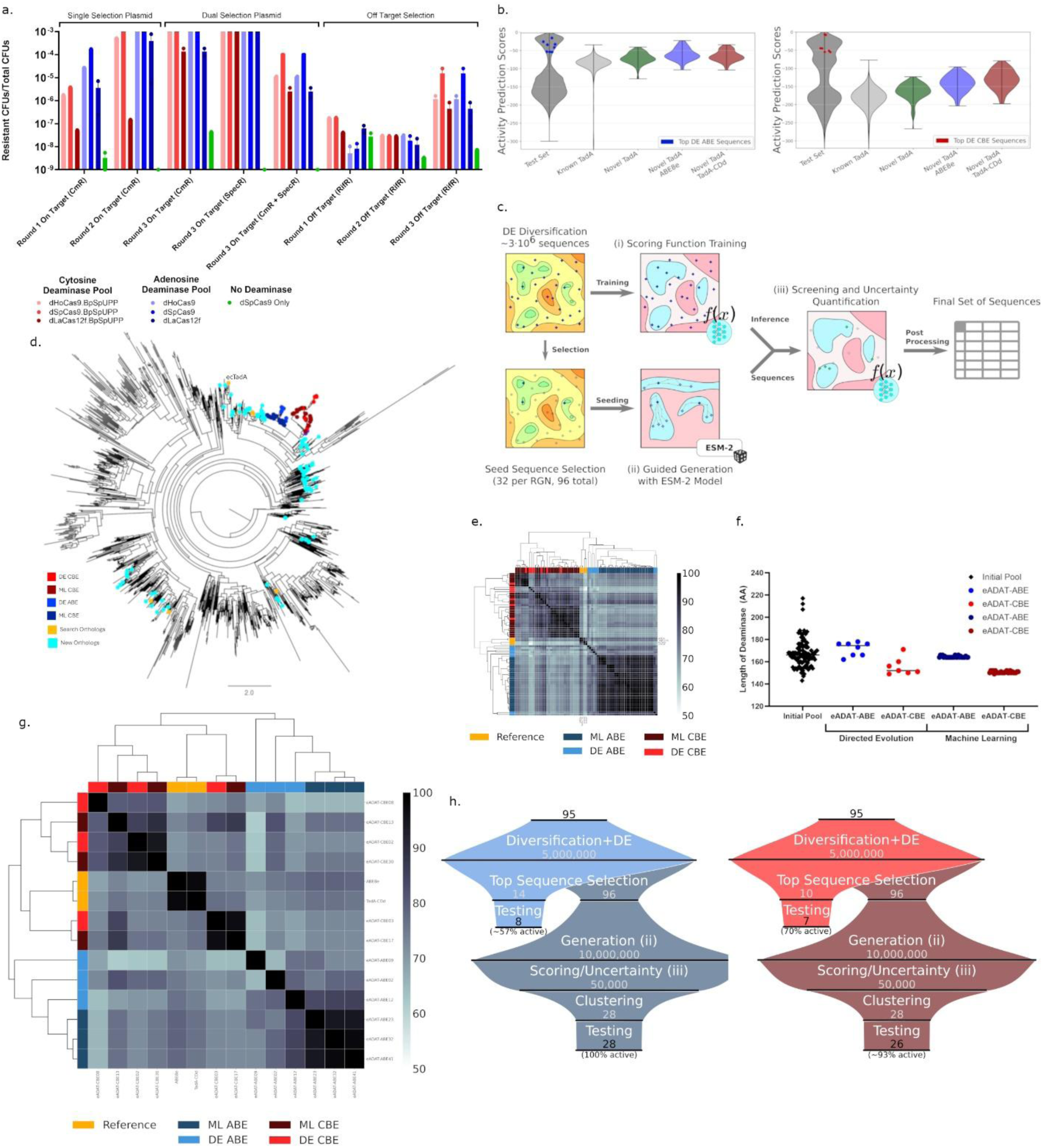
(a) Bacterial enrichment of active deaminases. Cultures were diluted to give accurate counts of selective vs. non-selective CFUs in a range of 10^-3^ to 10^-9^ from each successive round of enrichment. Each point represents an independent selection transformation, and the bar represents the average. Bars extending beyond the y-axis limits exceed the detection limit. (b) Machine Learning Model Validation. Predicted log score distributions using our scoring functions (ABE trained data left, CBE trained data right). Test Set indicates the holdout sequences from DE (dark grey). Known TadA Sequences are in light grey. Novel TadA corresponds to the 95 sequences used for initial diversification (Green). Novel TadA ABE8e (Blue) incorporate ABE beneficial mutations while Novel TadA-CDd (Red) incorporate CBE beneficial mutations in the original sequences. Top ABE/CBE sequences selected from DE that showed eukaryotic activity are red and blue dots accordingly. (c) Machine Learning data driven Virtual Screen. The scoring functions to learn the activity landscape were trained on the original sequences (i). Concurrently, the most active sequences were used as seed sequences for the ESM-2 iterative mask filling algorithm (ii), to introduce multiple co-dependent mutations while traversing the general protein fitness landscape. The generative algorithm (ii) gave rise to 10^7^ biologically plausible sequences which were filtered for their activity as base editors (iii). (d) Phylogenetic tree of ADAT family. Reference ADAT sequences are unmarked lines. Others are marked as indicated. (e) Percent identity comparison of deaminases derived from ML or DE. (f) Length of the deaminase proteins in amino acids. (g) Percent identity comparison of top eukaryotic performing deaminases derived from ML or DE. (h) Diversification and selection funnel for ABE (left) and CBE (right) for *in vitro* DE (lighter shade) and *in silico* ML (darker shade) screens. Successful eukaryotic activity is shown at the bottom of each.

Also of note, the rate of activity for dLaCas12f throughout all rounds—although elevated over the dSpCas9 control—was significantly lower than the activity rate observed in HoCas9 and SpCas9, potentially related to its lower activity overall in eukaryotes (p = 3.7e-7; Supplemental Table 3). All three rounds of enrichment were performed without additional diversification after initial DNA shuffling. Instead, all the surviving sequences were advanced to the next round. After each round, the deaminase pools were isolated and underwent next generation sequencing. By comparing the abundance of specific sequences across rounds, we were able to identify the top enriched sequences, which subsequently underwent validation in eukaryotes (Supplemental Tables 8&9).

### Machine Learning Scoring Function

We trained a range of deep neural networks to serve as scoring functions to optimize proteins for properties of interest. The model selection and evaluation were performed using 4-fold cross validation. Due to the imbalance of the classification task, the best models were selected based on the delta area under Precision Recall Curve (dAUPRC) which achieved scores of 0.30 (±0.03) and 0.25 (±0.02) on average for ABE and CBE, respectively. The selected models were tested on the holdout sets and reached an average area-under-receiver operating curve (AUC) of 0.96 (±0.01) and 0.93 (±0.01) for ABE and CBE respectively, indicating a high predictive quality of the models. These models were obtained by training in a multitask setting for the two base editors separately, with multiple enrichment thresholds and per-task class balancing.

To validate the resulting scoring functions, we visualized the predicted scores for various sequences, as well as those from the holdout test set (Figure 1b). The predicted scores are log probabilities of achieving at least 50x enrichment in the DE experiment. The holdout test set shows a bimodal distribution of sequences predicted to have either high or low activity. The middle region likely corresponds to sequences for which the models have low confidence, which could be a result of subtle structural differences placing them more out of distribution compared to our DE training dataset. The top adenosine deaminase and cytosine deaminase sequences from DE are ranked highly by their respective models. As expected, models trained on cytosine deaminases tend to rank the CBE-activating mutations higher than the ABE-activating mutations, and vice versa. Sequences with conserved mutations introduced from ABE8e or TadA-CDd were ranked higher than the wildtype sequences by both models, which could be explained by the mutations enabling base level of activity or by the high degree of overlapping mutations introduced into the ABE and CBE pools and subsequently introduced into the training set. The full family of known TadA orthologs(25) (Supplemental Table 10) tended to score lower than our 95 parent wildtype sequences, potentially due to greater sequence divergence from the training set. Based on the distribution of scores, known TadA orthologs are predicted to achieve lower performance as CBEs while, at the same time, the ABE models rank TadA orthologs between the two distribution modes. The low-activity group overlapped with the known TadA orthologs for the CBE predictions but showed noticeably lower scores than known TadA orthologs in the ABE predictions. It is reasonable that our model may have lower confidence in classifying native ADATs outside our training set as adenosine base editors because TadA orthologs are natively RNA adenosine deaminases, with only limited reports of ssDNA cytosine deamination by some TadA orthologs, and we did not fully explore the entire sequence space of the ADAT family.(79) The high predictive quality of the scoring functions thus made it possible to perform an *in silico* virtual screen of millions of novel sequences for advancement to *in vitro* testing.

### Machine Learning Sequence Generation

To generate novel sequences for the virtual screen (Figure 1c), the top 32 sequences with the highest enrichment score were selected for each of the three RGNs, for a total of 96 seed sequences for subsequent generative modeling using the ESM-2(31) protein language model (PLM). The 650-million parameter model from that model family was used to suggest beneficial mutations at various locations around the seed sequences for each base editor. A generative algorithm was required to productively narrow the search space, as exhaustive sampling is infeasible for more than a small number of mutations, even for virtual screening.(36) The generation algorithms based on protein language models are suited to dealing with variable-length sequences and large mutation distances that can result in pronounced epistasis effects.(80) Our iterative ESM mask-filling generation was used as the proposal algorithm (detailed in Methods), which allowed us to incorporate global protein fitness knowledge beyond the scope of the generated DE data into the generation process.(31,32,81) Additionally, this process allowed us to introduce substitutions without a mutational bias, enabling us to explore novel mutations that would not likely arise through a random mutation approach.(82) In this way, we generated two separate screening pools for ABEs and CBEs, each containing 10 million sequences with up to three mutations from the corresponding reference sequences (Supplemental Figure 7).

Our scoring functions were then used to predict the fitness of generated sequences (see above; detailed in Methods). In addition to the score, the uncertainty estimation algorithm provided a confidence value for each prediction. The scoring functions and the uncertainty algorithms were used to virtually screen a large set of candidate sequences. The sequences were selected to have the highest scores and low uncertainty in order to obtain sequences with high predictive activity and still be within the training domain of our dataset.^38, 44^ This combination of scoring and uncertainty estimation allowed us to select a set of promising sequences in a single step. The top 50,000 sequences by predicted ABE or CBE activity from the ML model were further filtered to prioritize 28 sequences for eukaryotic validation for each editing type (detailed in Methods). Those sequences were chosen to be consistently high scoring with low uncertainty, diverse in sequence, and contain mutations at, or near, regions homologous to those known to impact base editing in ecTadA (Supplemental Tables 8&9).

### Sequence Comparison of Selected Sequences

To compare the ML-suggested sequences to previously described TadA orthologs(25), we generated a phylogenetic tree (Figure 1d, Sequences listed in Supplemental Tables 5-11). Although we identified a diverse group of orthologs, they do not span the range of TadA sequence space but sample heavily from selected regions. This was likely caused by our initial selection of input sequences, which was necessary to maintain regions of microhomology for efficient DNA shuffling. Active sequences derived from both DE and ML co-locate to a specific region of the tree that is, on average, 58% identical to the initial ecTadA protein, despite being selected for independently. Additionally, the alignment of active sequences reveals that they cluster first by ABE or CBE, and that ML sequences demonstrate greater similarity than those sequences identified by DE (Figure 1e). The greater similarity within the ML sequence cluster could be due to a few seed sequences dominating the results of the virtual screen.

While both pools started with the same size range of initial orthologs (143-217 amino acids, with an average of 168), the resulting sequences differed markedly in length between ABE and CBE, but not between DE or ML (Figure 1f). ABE-selected sequences averaged 172 and 164 amino acids for DE and ML sequences, respectively, while CBE-selected sequences averaged 156 and 151 amino acids for DE and ML sequences, respectively, representing an 8% decrease in size relative to the ABE sequences. The size difference was determined to be caused by a shorter carboxy termini relative to ecTadA, and not deletions elsewhere in the sequence (Supplemental Figure 8), although some DE sequences did have an extension to their amino terminus. The shorter CBE constructs ended at the R152P mutation which was introduced early in the evolution of ABE7.10, or directly thereafter at R153.(3) Notably, the R152P mutation is known to be disruptive to the α5-helix and causes a sharp 180° turn at position 152 in ABE8e.(83) This shorter length was observed with multiple carboxy termini derived from independent sequences during DNA shuffling, despite the relative abundance of mutations introduced in this region during the evolution of ABE8e and TadA-CDd and the demonstrated importance of the combined R152P, E155V, I156F, K157N mutations in ABE8e.(83) In structures predicted using AlphaFold2(47), the same disruption of the α5-helix was observed in the adenosine deaminases as in ABE8e (Supplemental Figure 9).

### On-Target Eukaryotic Activity of Leading Deaminases

The top enriched sequences with high diversity (<85% (ABE) or <90% (CBE) homology) from DE were tested for eukaryotic activity across a limited number of targets and when fused to three different Cas9 enzymes, nSpCas9, nHoCas9, and nNme2Cas9, as well as BpSpUPP in the case of cytosine deaminases (Supplemental Figure 10).(51) Sequences showed higher activity when fused to nSpCas9 than the other two Cas9s tested, but all three RGNs matched on the relative activity of each deaminase. Overall, 8 of 14 ABE sequences and 7 of 10 CBE sequences derived from DE demonstrated some level of activity in eukaryotic cells. The top three performing sequences were selected for in-depth characterization across a range of eukaryotic targets. The top ML-generated sequences were tested on nSpCas9, and the top three performing sequences with collectively high diversity were also selected for in-depth characterization across a range of eukaryotic targets (Supplemental Figure 11). These top sequences show a range of diversity and cluster first by ABE or CBE activity (Figure 1g). However, while both ABE and CBE ML demonstrate greater sequence similarity than those sequences identified by DE, the diversity of ML CBE sequences is greater than ABE sequences. All 28 ABE sequences and 26 of 28 CBE sequences generated by ML, demonstrating a much higher success rate with sequences generated from our virtual screen than those generated by DE alone (Figure 1h).

When compared to ABE8e across eight different targets in eukaryotes, two of the three DE sequences had significantly lower activity than ABE8e (p < 0.05), whereas the remaining sequence and all three ML sequences were not significantly different from ABE8e (Figure 2a). When compared to TadA-CDd, all three DE sequences had significantly lower activity than TadA-CDd (p < 0.05), whereas all three ML sequences were not significantly different from TadA-CDd (Figure 2b). High off-base activity, i.e. cytosine deamination by an adenosine base editor and vice versa, was only observed with TadA-CDd but not with any other deaminases. All novel CBEs and three novel ABEs had a statistically significant reduction in off-base activity compared to the known base editors (p < 0.05) (Supplemental Figure 12). The lower off-base activity of two DE sequences, eADAT-ABE02 and eADAT-ABE12, may be attributable to their lower activity overall, but this explanation is insufficient to describe the lower off-base activity of the ML sequence, eADAT-ABE23, as it had on-target activity comparable to ABE8e. We observed no strong dinucleotide context preferences for the targeted adenosine or cytosine in DNA for any of our sequences, including control sequences (Supplemental Figure 13).

**Figure 2.**
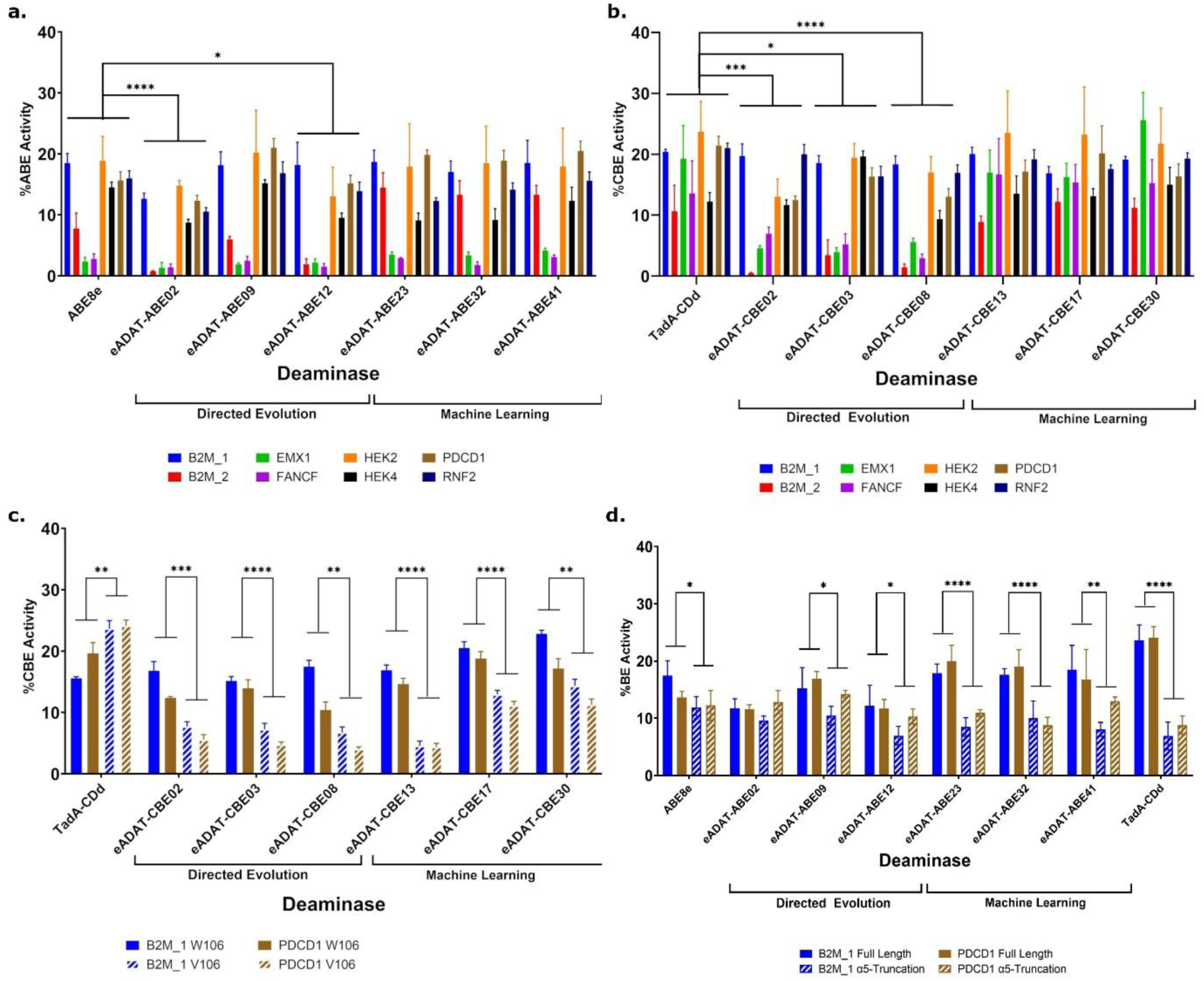
On Target Eukaryotic Validation of Top Deaminases. Mean of the replicates is shown and Error bars represent the standard error of the mean. *=p<0.05, **=p<0.005, ***=p<0.0005, ****=p<0.00005 (a) Top adenosine deaminases across eight targets. n=3 replicates for experimental conditions and n=5 replicates for controls. Statistically significant differences from ABE8e are indicated, and all remaining comparisons to ABE8e showed no statistically significant difference. (b) Top cytosine deaminases across eight targets. n=3 replicates for experimental conditions and n=5 replicates for controls. Statistically significant differences from TadA-CDd are indicated, and all remaining comparisons to TadA-CDd showed no statistically significant difference. (c) Top cytosine deaminases with and without the V106W TadA-CD specificity mutation across two targets. n=3-4 replicates. Statistically significant differences between the Valine or Tryptophan at homologous residue to ecTadA 106 are indicated. (d) Top adenosine deaminases with α5-helix truncations across two targets. n=3-4 replicates. Statistically significant differences between the full length or truncated deaminase are indicated. Remaining comparisons showed no statistically significant difference.

The V106W mutation in TadA, which is known to improve C•G-to-T•A selectivity in TadA-CDd(21), was introduced into the original pool for cytosine deamination activity and was present in all selected sequences. To investigate its effect, we compared variants with equivalent W106V mutations to TadA-CDd with and without V106W. As previously reported, we observed a decrease in activity when V106W was incorporated to TadA-CDd.(21) However, when we back-mutated the homologous V106W mutation in our new sequences, we instead observed that tryptophan outperformed the valine residue in on-target activity (Figure 2c). Additionally, we observed no change in off-base activity between our sequences containing valine or tryptophan at the homologous position, but we did observe the expected decrease in off-base activity for TadA-CDd_V106W (Supplemental Figure 14). There were no significant differences between any of our six original sequences and TadA-CDd_V106W in on-target or off-base activity. This suggests that the reported impacts of the V106W mutation may be specific to ecTadA, or that our selection methods successfully enriched for unknown compensatory mutations in our novel backgrounds.

### Effects of C-Terminus Truncation on Activity

It was previously shown that elongating the linker between the C-terminus of the deaminase and Cas9 in BE3 from 16 amino acids to 32 amino acids did not change the editing window, but did increase the editing rate, which was hypothesized to occur by enhancing access to the ssDNA in the R-loop formed by Cas9.(4) Here, however, we see comparable editing with our eADAT-CBE constructs compared to TadA-CDd, despite the shorter C terminal domain(Figure 2b).

The editing window was largely unchanged for the novel, active deaminases compared to ABE8e and TadA-CDd, from positions 2 to 11 in the target sequence for the ABEs and from 2 to 9 for the CBEs, with activity peaking at positions 3 to 8 for both (Supplemental Figure 15). This was despite the absence of approximately 14 amino acids at the carboxy end of the cytosine deaminases compared to TadA-CDd.

To determine if smaller adenosine deaminases would perform equally well, despite the observed DE preference for a longer sequence, we truncated all sequences after the residue homologous to TadA R153 based on a multiple sequence alignment in our ABE sequences. Both ABE8e and TadA-CDd showed a decrease in activity from the deletion of 14 residues from the α5-helix, as did most of our ABE constructs, with the ML construct showing more deleterious effects than the DE constructs (Figure 2d). This requirement for an elongated C terminus in ABE deaminases suggests that our model is pulling out a bonified requirement for a longer, but disrupted, α5-helix in ABE but that it is truly not required for active CBE deaminases.

### Cas-Independent Off-Target Editing of Leading Deaminases

Deaminases with high activity also tend to have undesired Cas-independent off-target editing, which can prevent their use in some applications.(9,11,13–16) ABE8e and TadA-CDd both have minimal Cas-independent DNA off-target editing compared to eukaryotic deaminases.(21) To confirm that our newly developed deaminases maintain this feature, we measured their off-target activity with an orthogonal R-loop assay.(13) In this assay a hotspot of single-stranded DNA is created by a gRNA-targeted dead Cas9 enzyme, while a base editing construct is simultaneously delivered and targeted to a distinct and separate location in the genome, allowing quantification of potential Cas-independent off-target activity of the base editor. We compared the performance of our deaminases against three independent sites in the human genome (Figure 3a&b). We included a base editor developed with the human APOBEC3A (hA3A) protein, a well-known eukaryotic cytosine deaminase with high on and off target activity as a base editor.(16) For ADAT derived ABEs and CBEs we saw no increase in Cas-independent off-target editing compared to the respective control, and saw the expected high off target activity with hA3A. The on-target editing again showed that some DE sequences had lower on-target editing than ML sequences, with eADAT-ABE2 significantly lower than ABE8e (p=0.0038) and eADAT-CBE02, eADAT-CBE08, and eADAT-CBE13 significantly lower than TadA-CDd (p=0.0015, 0.0013, 0.0094 respectively) (Supplemental Figure 16).

**Figure 3.**
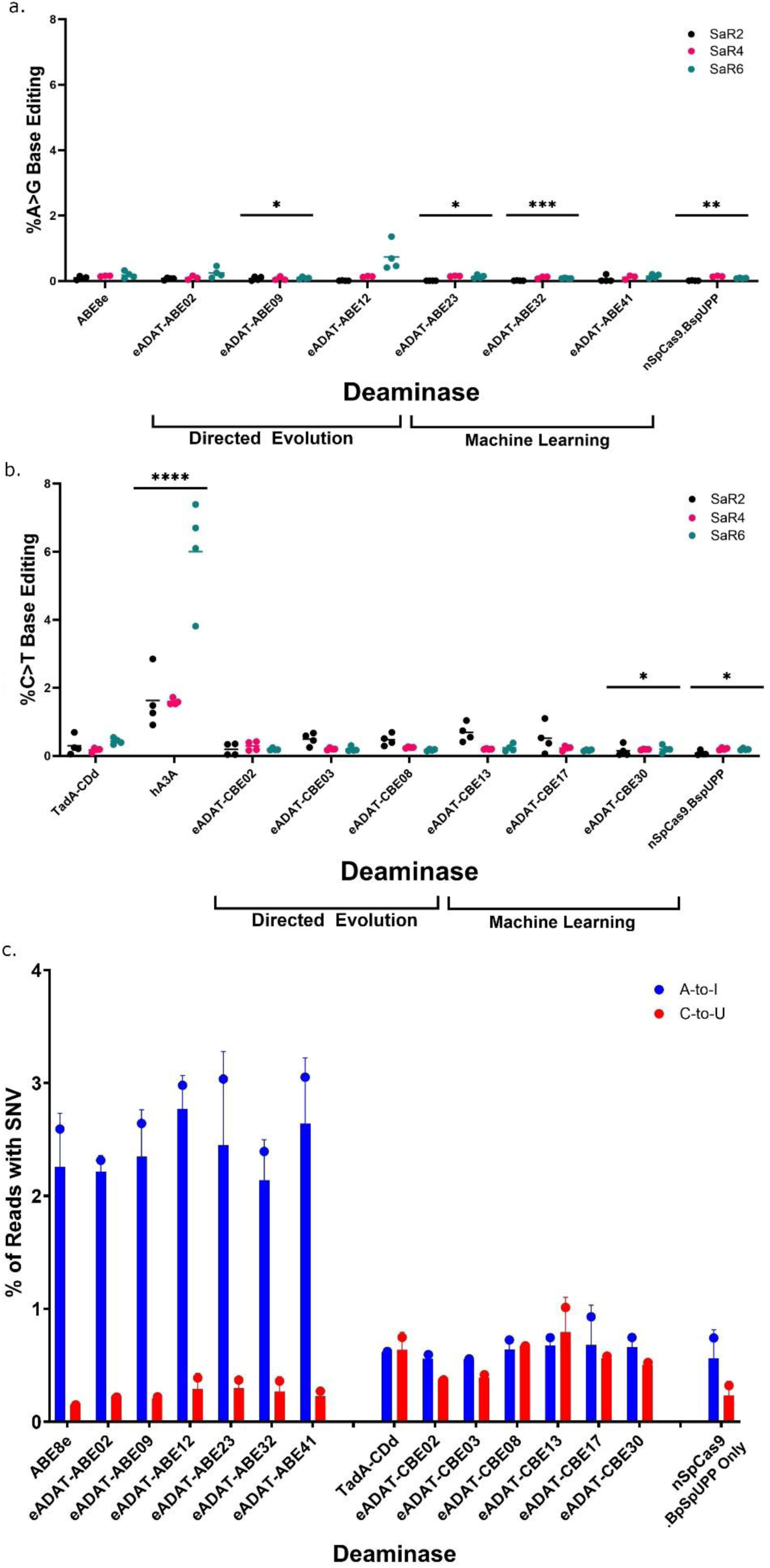
Off Target Eukaryotic Validation of Top Deaminases. Off Target DNA Base editing activity of leading adenosine deaminases (a) or cytosine (b) at off-target DNA sites created by an orthogonal Cas9 enzyme. Lines indicate the mean of 3-4 replicates. *=p<0.05, **=p<0.005, ***=p<0.0005, ****=p<0.00005 relative to respective control. (c) Cas-independent off-target RNA editing of leading deaminases. Bars represent the mean of n=2 replicates.

Our deaminases were also evaluated for off-target RNA editing by measuring the rate of SNV formation in the transcriptome. Because TadA natively functions on RNA, the overexpression of ABE8e has been shown to increase SNVs in the transcriptome.(84) We overexpressed our constructs and quantified the novel SNVs in the transcriptome compared to a non-transduced control (Figure 3c, and Supplemental Table 12). All the adenosine deaminase constructs showed low but detectable increases in A-to-I editing but not C-to-U editing compared to a nickase-only version of SpCas9; whereas some of the cytosine deaminase constructs, including TadA-CDd, showed a slight increase in C-to-U, but not A-to-I, editing. Off-target RNA editing was not significantly higher in the novel editors compared to the known editors. As with DNA editing, there was no strong preference for any dinucleotide by any of our sequences, including the control sequences. (Supplemental Figure 17).

## 4 Discussion

Base editing has shown tremendous promise as a therapeutic, with several clinical trials underway (National Library of Medicine [NLM], NCT05398029, NCT05456880, NCT05885464).(84) The directed evolution of ecTadA into variants that specifically catalyze either adenosine or cytosine offered improvements over the original base editor designs, which used eukaryotic-derived cytosine deaminases.(1,3,21) However, relatively little effort has been spent exploring alternative TadA orthologs compared to eukaryotic deaminases,(25) likely due to the challenge of evolving a new TadA ortholog to function with high activity. To overcome this, we developed a ML virtual screen which utilized a set of deep neural networks to design high-activity adenine and cytosine deaminases trained on experimental data from a large pool of novel TadA orthologs with only a single round of diversification. The scoring functions of our ML approach had a high predictive quality, comparable to a biological test, and could be viewed as a virtual screen.(85) By utilizing uncertainty estimation techniques(72,73) for the scoring functions we ensured that this *in silico* screen stayed within the domain of the training set and maintained a high predictive value for the resulting sequences.

This virtual screen improved upon a standard DE screen in several ways. First, the computational prediction can be done quickly, dramatically reducing the time and resources required to perform a screen, while still yielding highly active sequences with desired biological properties that were not present in the initial diversification pool.(85)

Second, the exploration of the sequence space in a typical DE process is random but also biased towards mutationally probably codons.(82) In contrast, by combining pretrained generative models with the DE dataset our generative modeling approach will only suggest mutations that are meaningful in the context of a given sequence. Our iterative mask filling algorithm utilized ESM-2 PLM and allowed us to obtain many high-quality candidate sequences compared to GPT-style PLMs, as well as sampling marginals of residues per position. Concurrent works(67,86) involving mask filling focus on searching for the best substitutions directly using the PLM as well as performing one substitution at a time. Instead, we sample the predictive distribution of multiple residues iteratively to produce a large volume of biologically plausible sequences, delegating the domain-specific selection to our scoring functions while not being constrained by the existing nucleotide composition of the seed sequence to only sample mutationally likely codons. Our approach could further benefit from utilizing more recent and capable PLMs such as ESM-3 or Bio-xLSTM.(30,32)

Third, our modelling approach can also be used to generate new rounds of sequences for further engineering efforts to improve additional properties, such as optimizing for differing dinucleotide context specificity or decreased cellular RNA editing. Bayesian uncertainty measures can be interpreted as prediction risk(87) with respect to specific modelling objectives.(88) Those measures, grounded in information theory, were used successfully to increase the rate of sequences that have shown activity in eukaryote validation compared to the DE alone approach. (87) The exponential decline in the proportion of variants that retain function as more random amino acids are introduced is a commonly encountered challenge in protein design, due to the chance that a random substitution will abolish protein function.(35) Our ML approach enables a prescreen for potentially active sequences with multiple mutations, allowing for the exploration of a larger protein landscape than random mutation, therefore minimizing the effort spent on sequences that lack support in the dataset and could as well have been sampled randomly. Extending the training set with meaningfully different sequences could further increase the predictive power and generalizability of the scoring models.

This approach allowed us to rapidly predict and test novel variants with comparable activity to the leading ecTadA-derived base editors. These novel editors are highly divergent from those derived from ecTadA and from each other, dramatically opening the space for further optimization efforts, while avoiding epistasis caused by using a single enzyme as an engineering starting point. In addition, the sequences derived from ML generally outperformed those selected by traditional enrichment, demonstrating the benefit of incorporating ML into protein engineering to develop high-activity enzymes with limited diversification and selection.

Notably, we observed that the discovered cytosine deaminases are shorter than the adenosine deaminases. Minimally engineered orthologs of TadA have been reported to function as ABEs, CBEs, and ACBEs but ranged in size from 152 to 350 amino acids.(25) This compact size did not derive from our initial diversification pool which had a uniform distribution of sizes for our initial sequences for both ABE and CBE. Additionally, the shorter length was observed across a range of different carboxy tails, which all removed the mutations introduced at the end of the α5-helix in the creation of ABE8e and TadA-CDd.(83) It is possible that the evolved CBEs do not need the same access to the non-template strand that the sharp 180° turn caused by the R152P substitution in ecTadA is proposed to enable, or that the longer α5-helix stabilizes the interaction with the larger purine base in adenosine deaminases. When our adenosine deaminases and ecTadA-derived deaminases were truncated to the same length, we saw that the truncated versions were generally less active than the original sequences, including TadA-CDd and ABE8e. The requirement for a longer α5-helix in TadA-CDd could be a result from its origins starting with ABE8e, and that additional mutations in the longer tail do play a role in its activity. Nevertheless we demonstrate that starting with a diverse pool of sequences preferentially results in smaller cytosine deaminases compared to adenosine deaminases. These new cytosine deaminases are now only 66% of the length of APOBEC1 used in BE1 and 90% of ecTadA, representing a marked decrease in size for CBEs.(1) This reduced size could also enable different attachment strategies, such as alternative internal fusions or linkers, which have been shown to be a successful avenue for improving the desired properties of base editors.(7)

There was noticeable improvement in off-base activity for both types of editors compared to reported editors, and off target RNA and DNA editing was comparable to reported editors. While our initial diversification pool incorporated known mutations that turned ecTadA into ABE8e or TadA-CDd, we also included V106W in the CBE pool. V106W is known to improve specificity of TadA-CDd for cytosine over adenosine, but reduces on-target activity.(21) It was possible that this mutation was responsible for the improved specificity of our CBE editors, although it was not incorporated into our improved ABE deaminases. However, when we removed the V106W mutation in our CBEs, we instead saw a decrease in on-target activity and no change in off-base activity, which is opposite to the effect seen in TadA-CDd. Unlike the original A106V and D108N mutations which first enabled ssDNA activity in the development of ABE7.10 and have been shown to enable ssDNA editing in novel TadA orthologs, the effect of the V106W mutation might be specific to ecTadA and could represent a potential effect caused by epistasis relying on a single starting sequence for deaminase engineering.(3,25)

While other studies have reported low, but detectable, Cas-independent DNA off-target editing via the R-loop assay we did not observe any such activity with any of the ADAT-derived deaminases.(17,27) We did observe Cas-independent off target with the eukaryotic hA3A deaminase as expected.(16) This could be due to differences in expression levels, caused by different precise promoter sequences or codon optimization, or by differences in experimental design, such as target selection, assay duration, or reagent concentration.(50) Because of this, we can not determine if our new designs outperform ABE8e or TadA-CDd in Cas-independent DNA off target activity, but we expect that they will at least perform comparably. Nevertheless, future studies into Cas-independent activity of ADAT derived deaminases could be aided by the demonstrated ability of our platform to develop highly active designs for investigation.

We had hypothesized that performing DNA shuffling across diverse orthologs and enriching for constructs with high DNA editing and no selection for RNA activity, would generate variants that created minimal RNA editing. However, RNA editing—although low—was not reduced compared to ABE8e or TadA-CDd. It may be that DNA and RNA activity are intrinsically linked and that only precisely introduced mutations allow fine discrimination between the two nucleic acid types, such as the original inclusion of V106W through careful examination of the ecTadA catalytic site.(54) Our approach could be used to accelerate investigations into this link by generating sequences with mutations at relevant positions, while still maintaining high predicted activity enabled by our virtual screen.

This study opens the space for new variants of base editors derived from TadA orthologs and demonstrates the benefits of incorporating ML and Bayesian deep learning methods into protein engineering efforts. This approach created multiple highly active diverse sequences that can be used for improved base editing applications, as well as for generating novel active sequences for future study.

## Acknowledgments

The metagenomic data were produced by the U.S. Department of Energy Joint Genome Institute (https://ror.org/04xm1d337; operated under Contract No. DE-AC02-05CH11231) in collaboration with the user community.

## Author contributions

MW, SM, and TB planned, executed and performed the wet-lab experiments. MI, JS, AM, GK developed the machine learning approach. MI, JS, and AM performed computational experiments including in-silico screening. LR and TB planned, implemented, and performed the search for novel orthologs and Cas enzymes. LS and LR performed the bioinformatics processing of the data. All authors analyzed and interpreted the results of the wet-lab and computational experiments. MI, MW, AM, JS, IMD, GK, and TB wrote the manuscript with input and revisions from all authors. IMD, GK, and TB conceived and designed the study. All authors have reviewed and approved the final version of the manuscript.

## Supplementary Data

Supplementary Figures: see after literature references

Supplementary Tables: File Supplementary_Info.xlsx provided; contains Supplementary Tables, where Tables 1-12 list amino acid sequences, gRNA, and coding sequences of samples used in this study as well as eukaryotic editing results of novel Cas12f enzymes

Supplementary Code: File acbe_minimum_code.pdf provided; contains minimum code required to reproduce computational experiments

## Conflict of Interest

MW, LS, LR, SM, IM, and TB are/were employees of UCB, receive salary from the company, and might own equity in the company. A patent application related to the article has been filed.

## Funding

This study was funded by UCB

## Data Availability

The datasets supporting the conclusions of this article are included within the article and its additional files. The training data of this study are available from the corresponding author, TB upon reasonable request.

## Supplementary Figures

We provide them at the following pages.

**Supplemental Figure 1.**
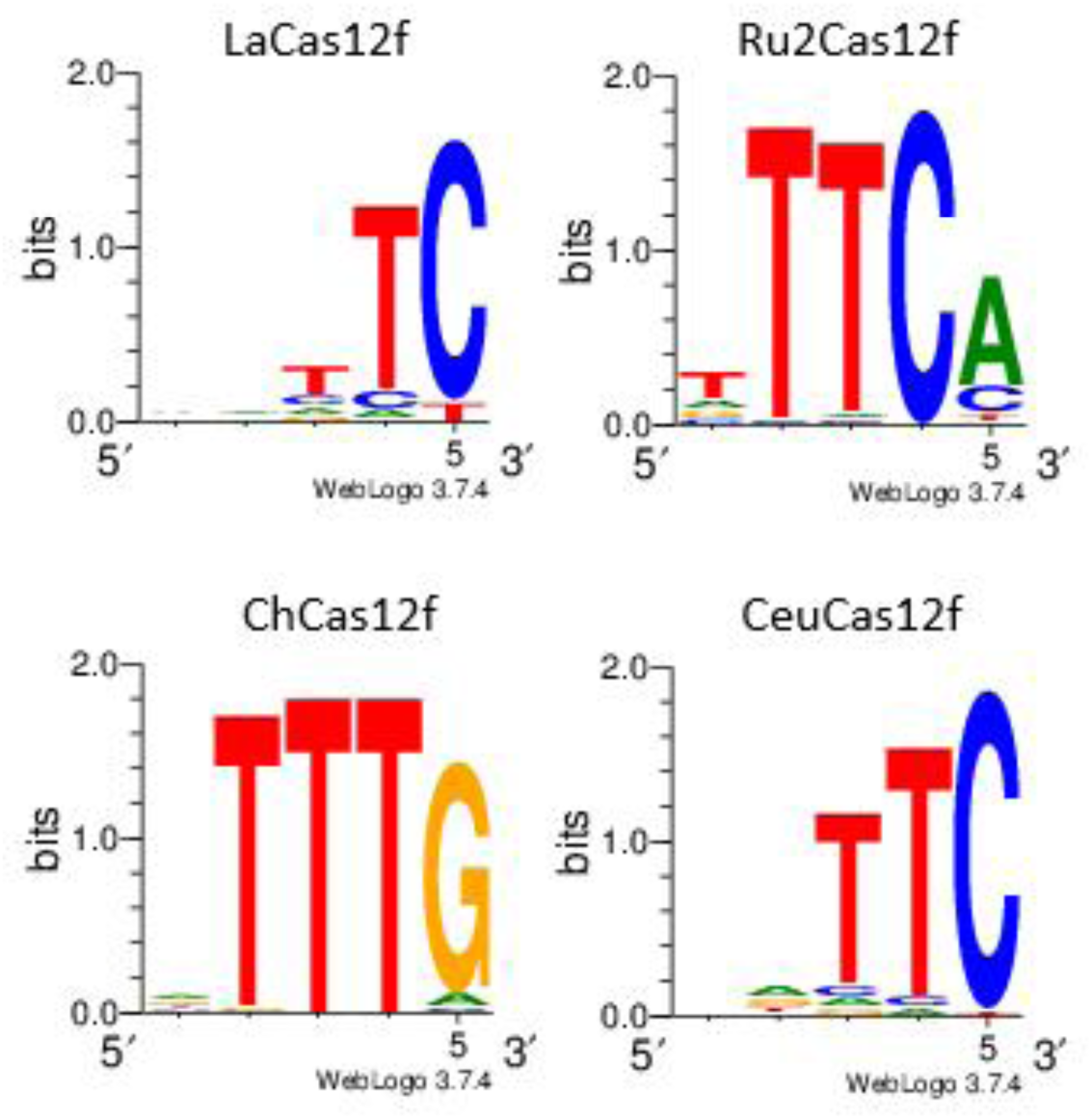
Identified 5’ PAM sequences of active Type V-F systems

**Supplemental Figure 2.**
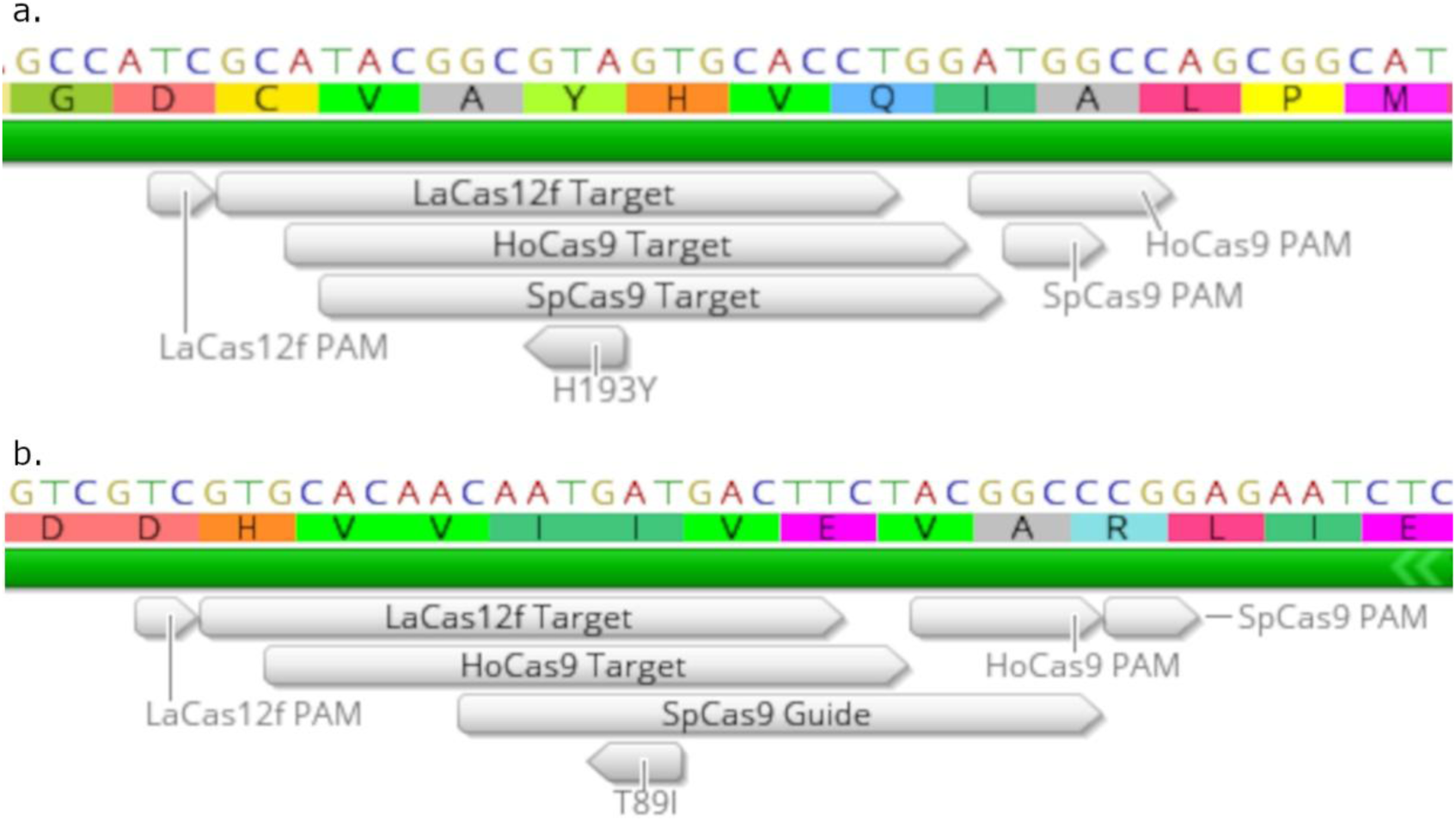
Target site for deamination to restore activity to dead Chloramphenicol (a) or Spectinomycin (b) resistance genes using adenosine base editors. Target sequence and PAM for each of the three RGNs used in the bacterial assay are indicated. Inactivating mutation capable of being corrected by an ABE indicated.

**Supplemental Figure 3.**
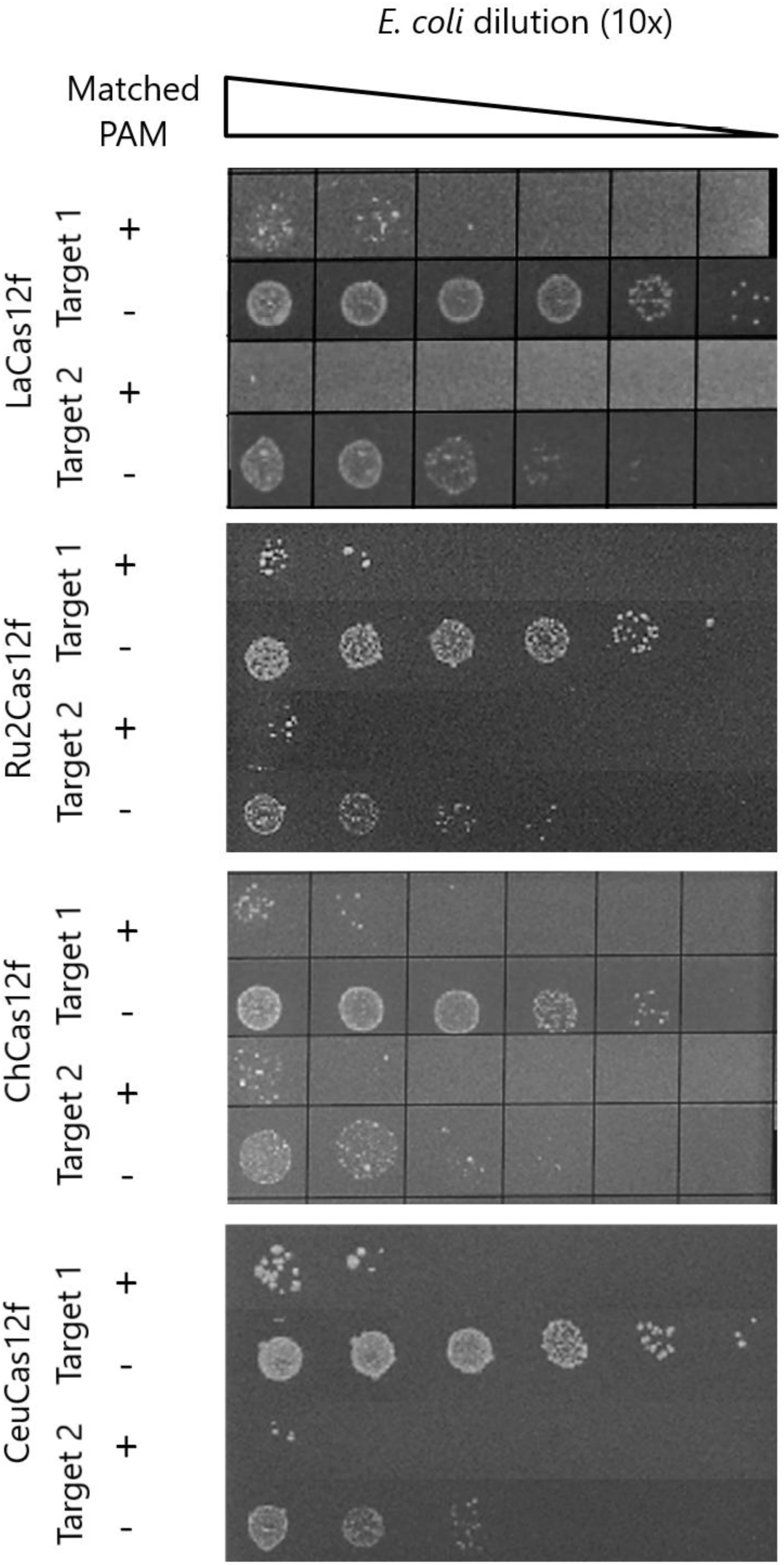
Cas12f mediated plasmid DNA interference in *E. coli*. Each system was compared with two different target sequences against matching target sequences, but with incompatible PAM sequences

**Supplemental Figure 4.**
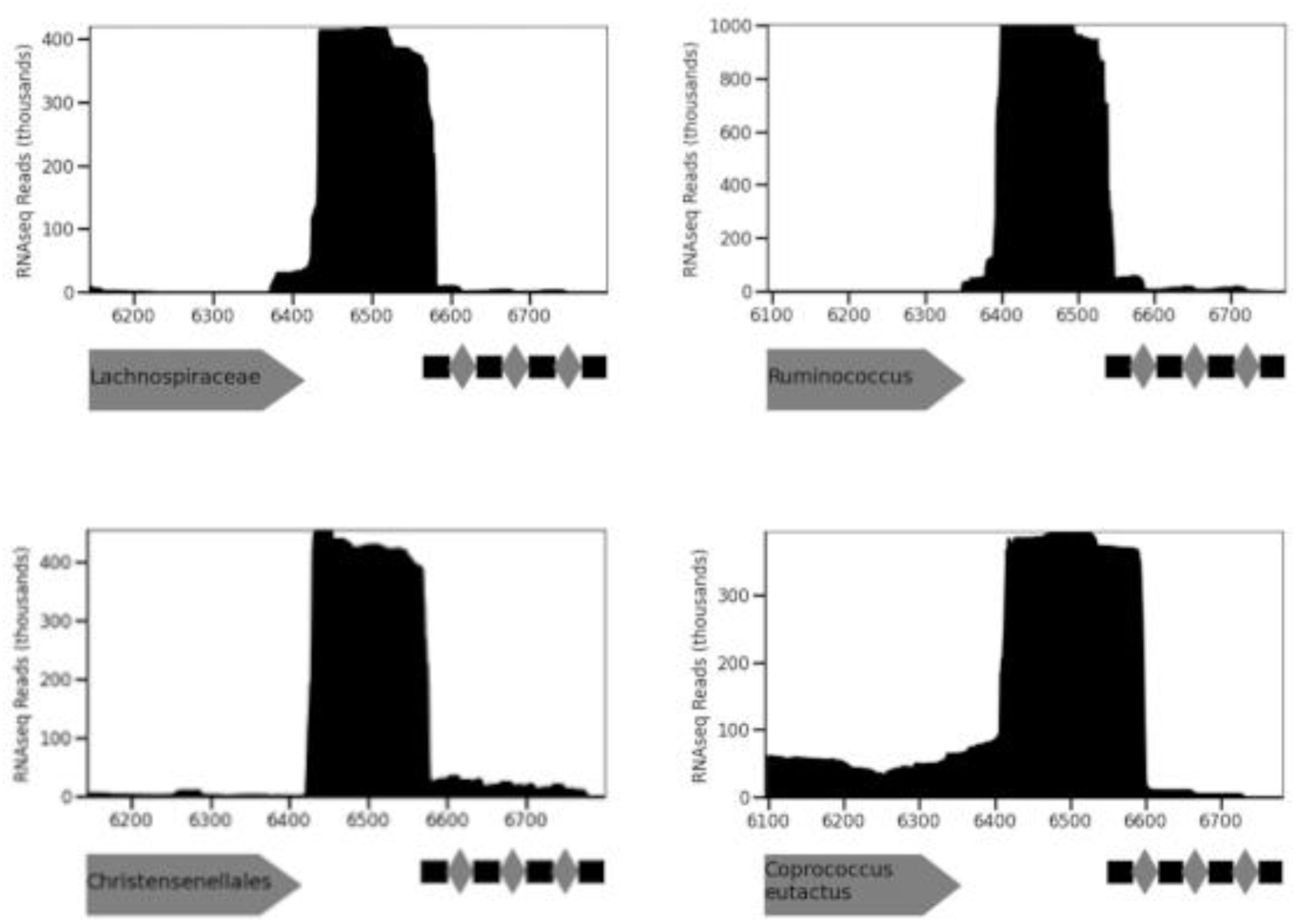
Small RNAseq of CRISPR operon of active Type Vf systems. Cas12f ORF is marked with a grey box with direction indicated by pointed edge. CRISPR array indicated with black boxes (repeats) and grey diamonds (spacers).

**Supplemental Figure 5.**
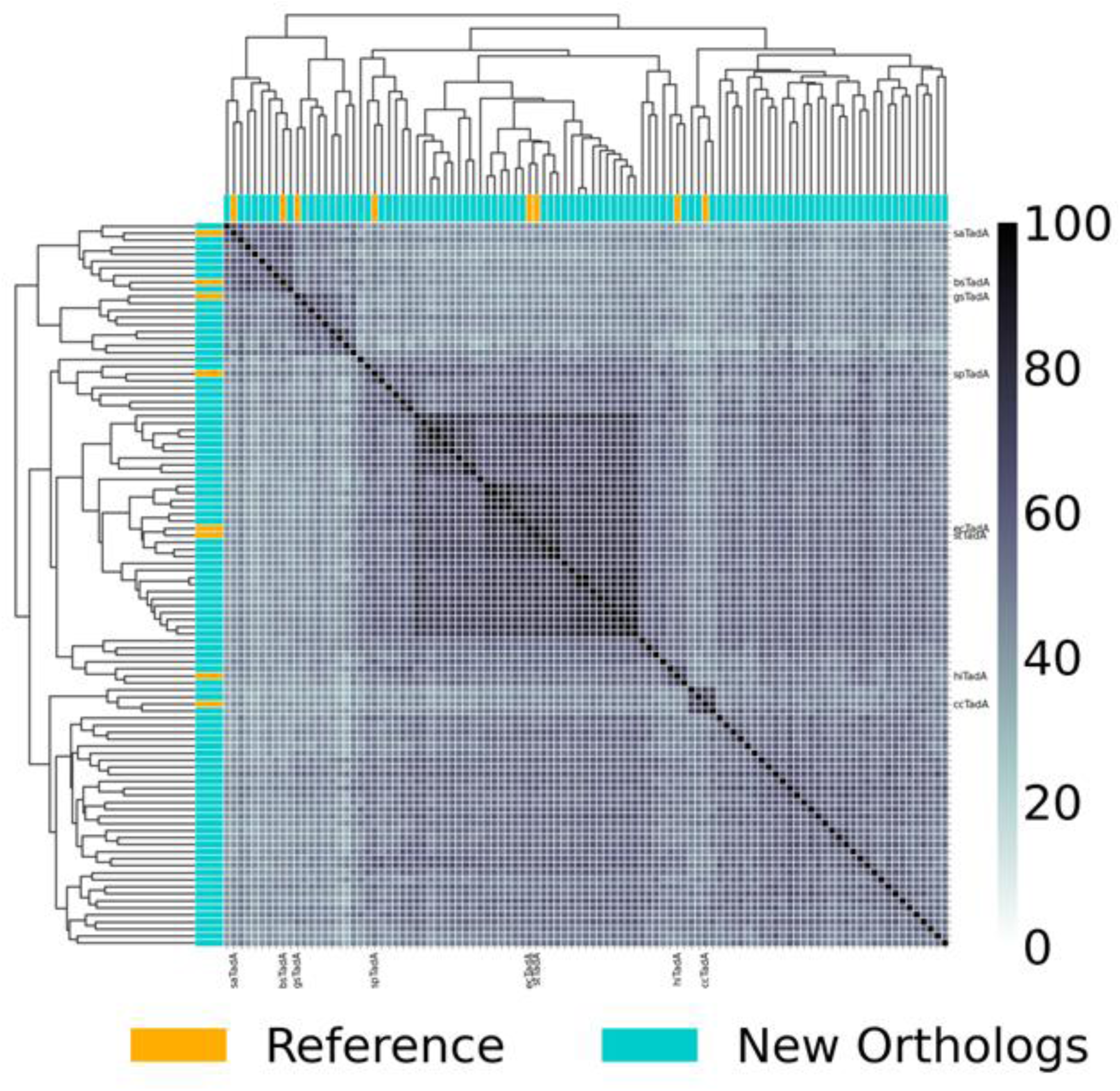
Heat map comparison of the percent identity of newly identified TadA orthologs compared to the original eight search sequences.

**Supplemental Figure 6.**
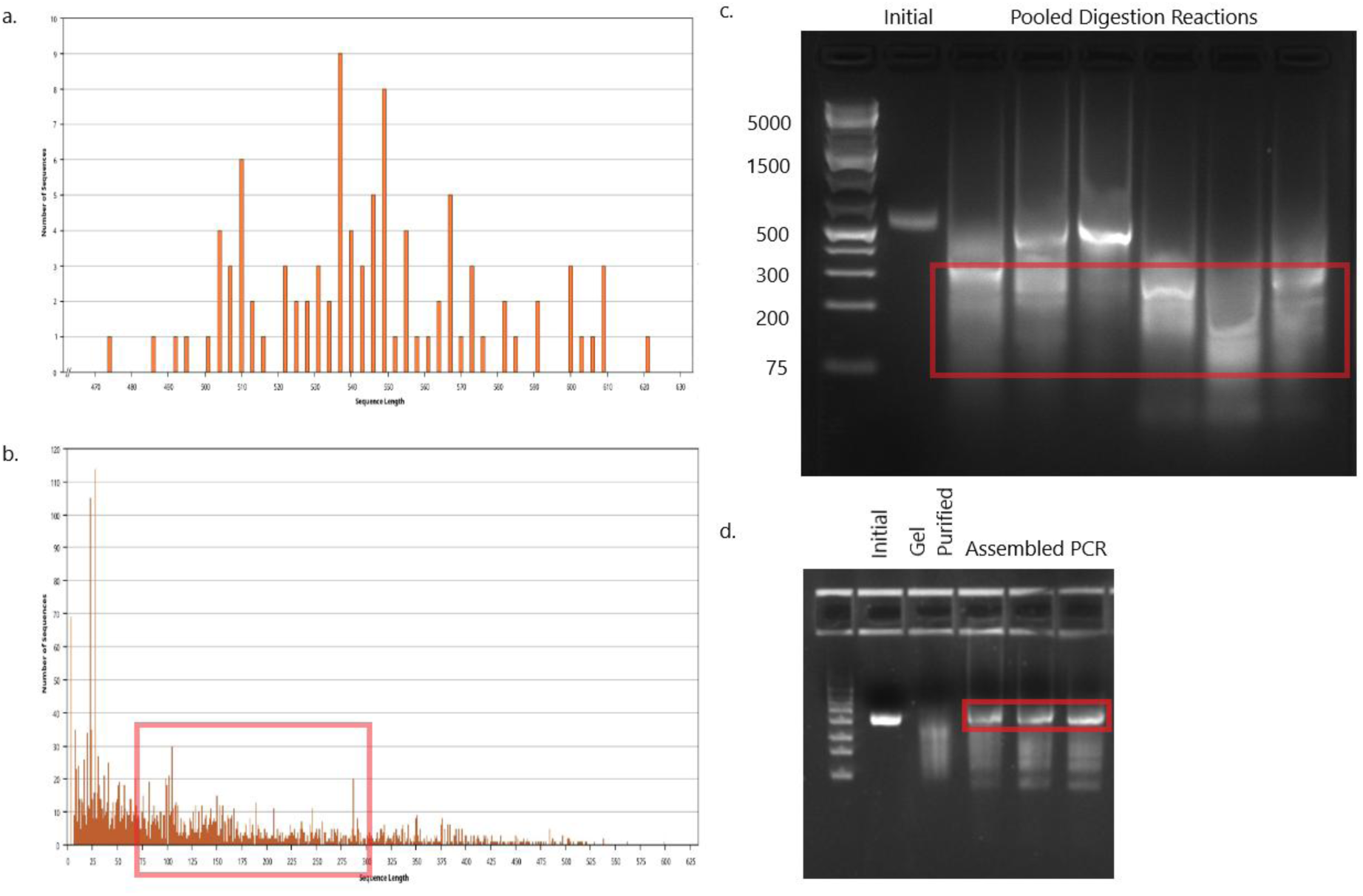
DNA Shuffling Assembly. The distribution of sequence lengths prior to digestion (a) and post digestion (b). The digested fragments (c) and the final amplified product (d). Red boxes indicate the region excised for selection into the next step

**Supplemental Figure 7.**
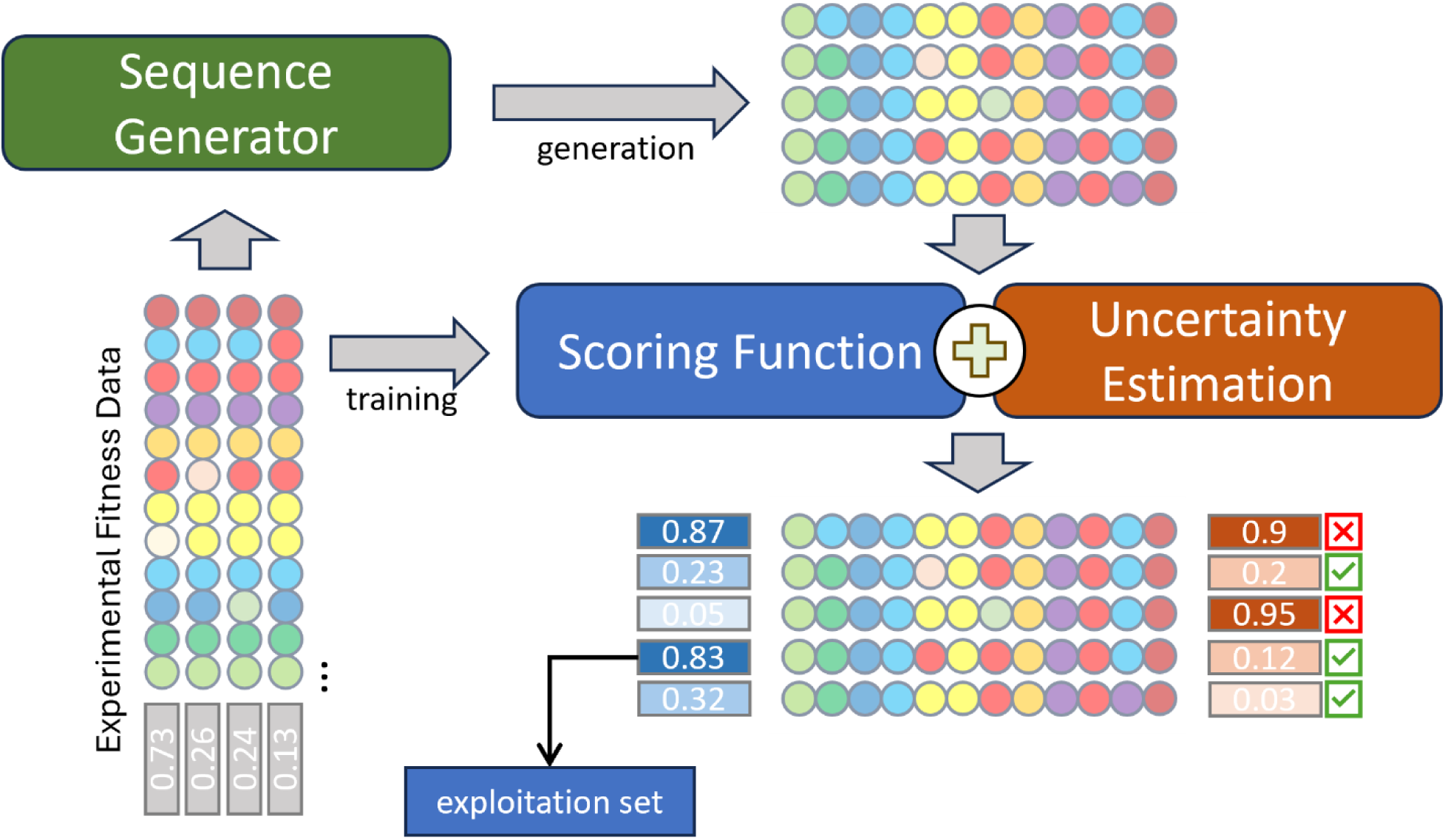
Overview of the machine learning approach combining a generative model with a scoring function. The experimental fitness data was obtained from the DE experiment measurements. The generative model samples restricted the considered distribution of sequences to that of feasible protein sequences based on the experimental data. The scoring function was trained on the experimental data and was used to rank the sequences sampled by the generative model. Uncertainty estimation provided the confidence values for the fitness estimates produced by the scoring function. The sequences with high predicted activity and high confidence were selected for experimental evaluation.

**Supplemental Figure 8.**
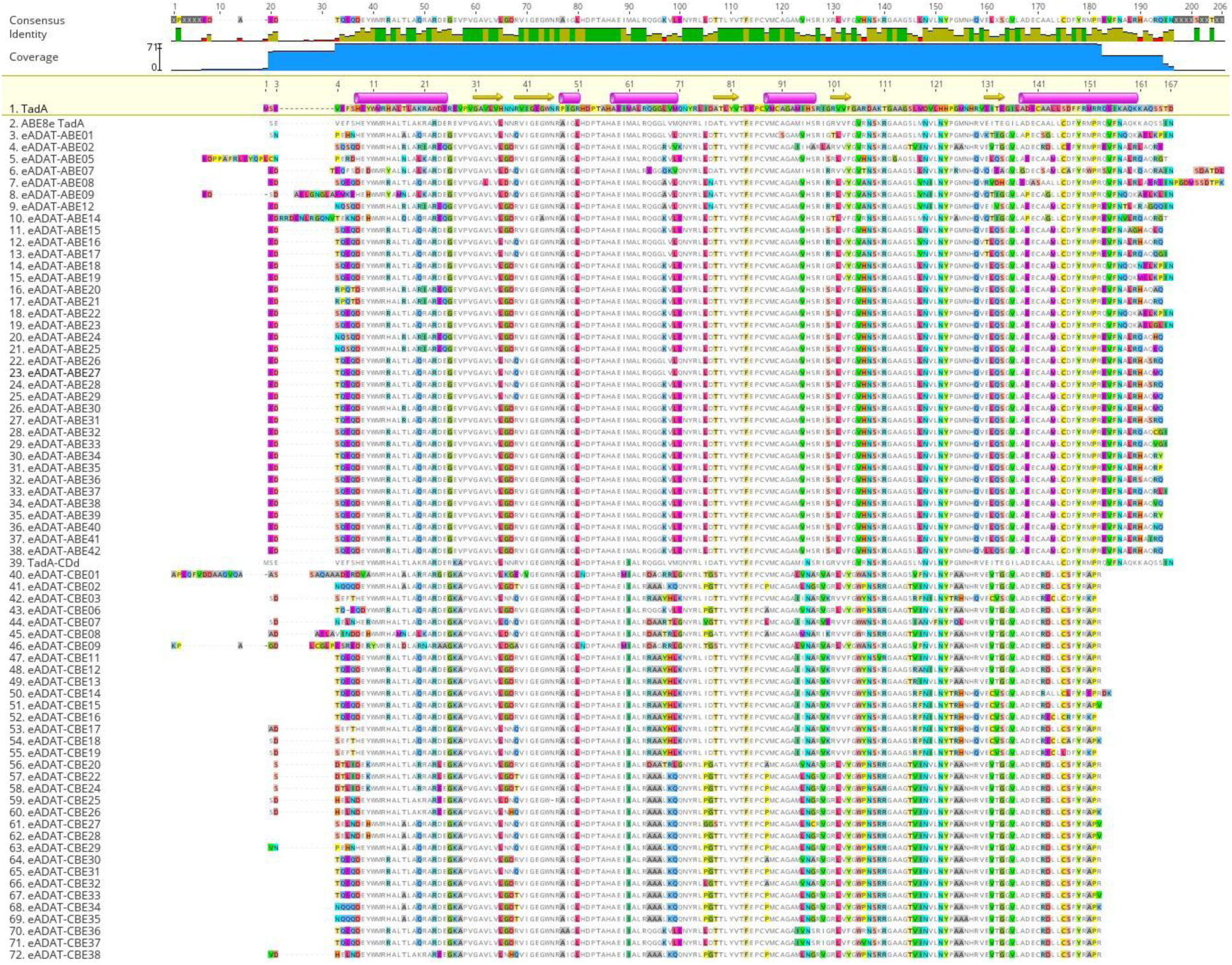
MAFFT E-INS-I Alignment of active sequences in eukaryotes to ecTadA. ABE sequences are on the top and CBE sequences are on the bottom.

**Supplemental Figure 9.**
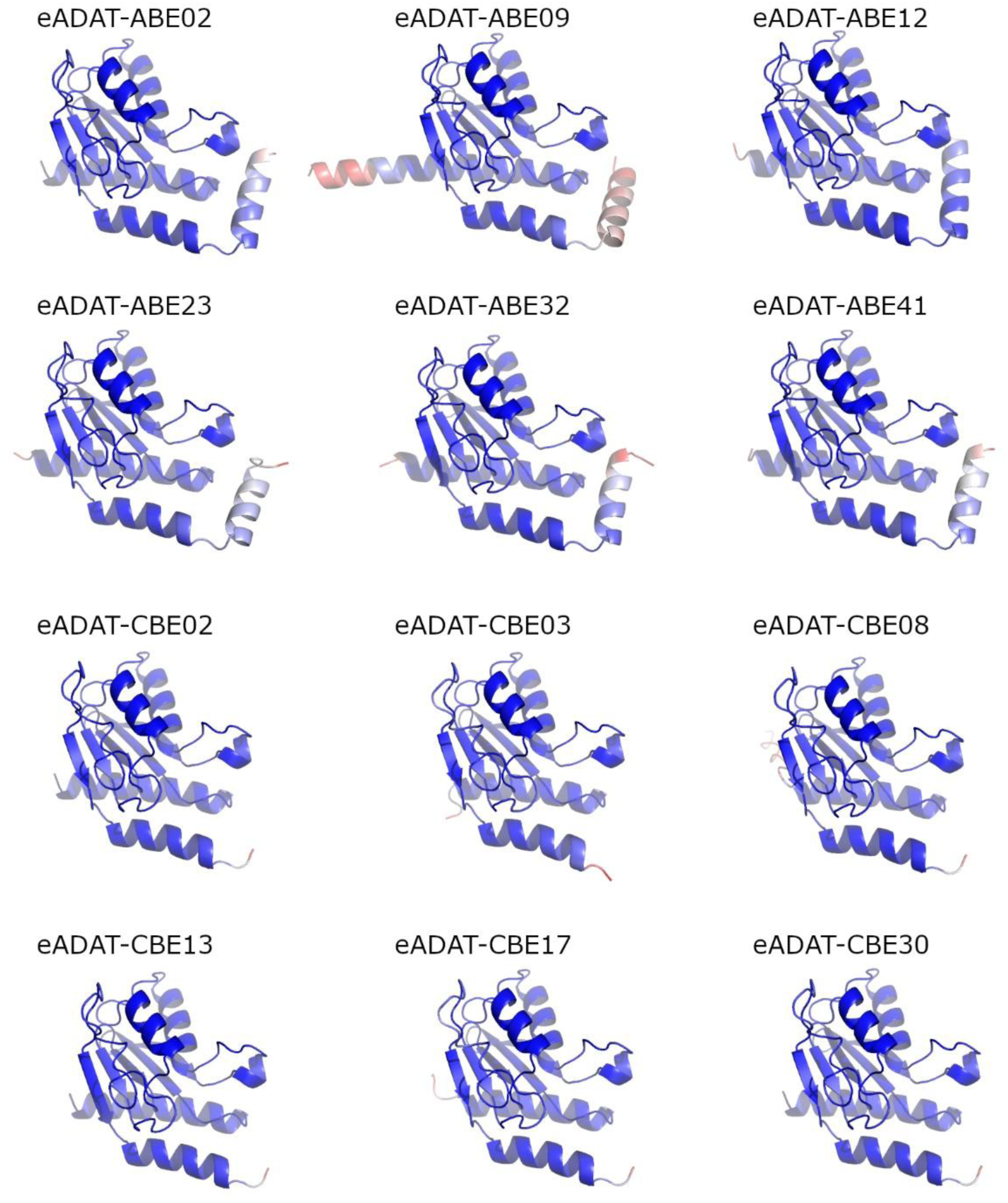
Predicted Alphafold2 structures of top 6 ABE (top) and top 6 CBE (bottom) deaminases. The structures are shaded by pLDDT values from 50 (red) to 100 (blue).

**Supplemental Figure 10.**
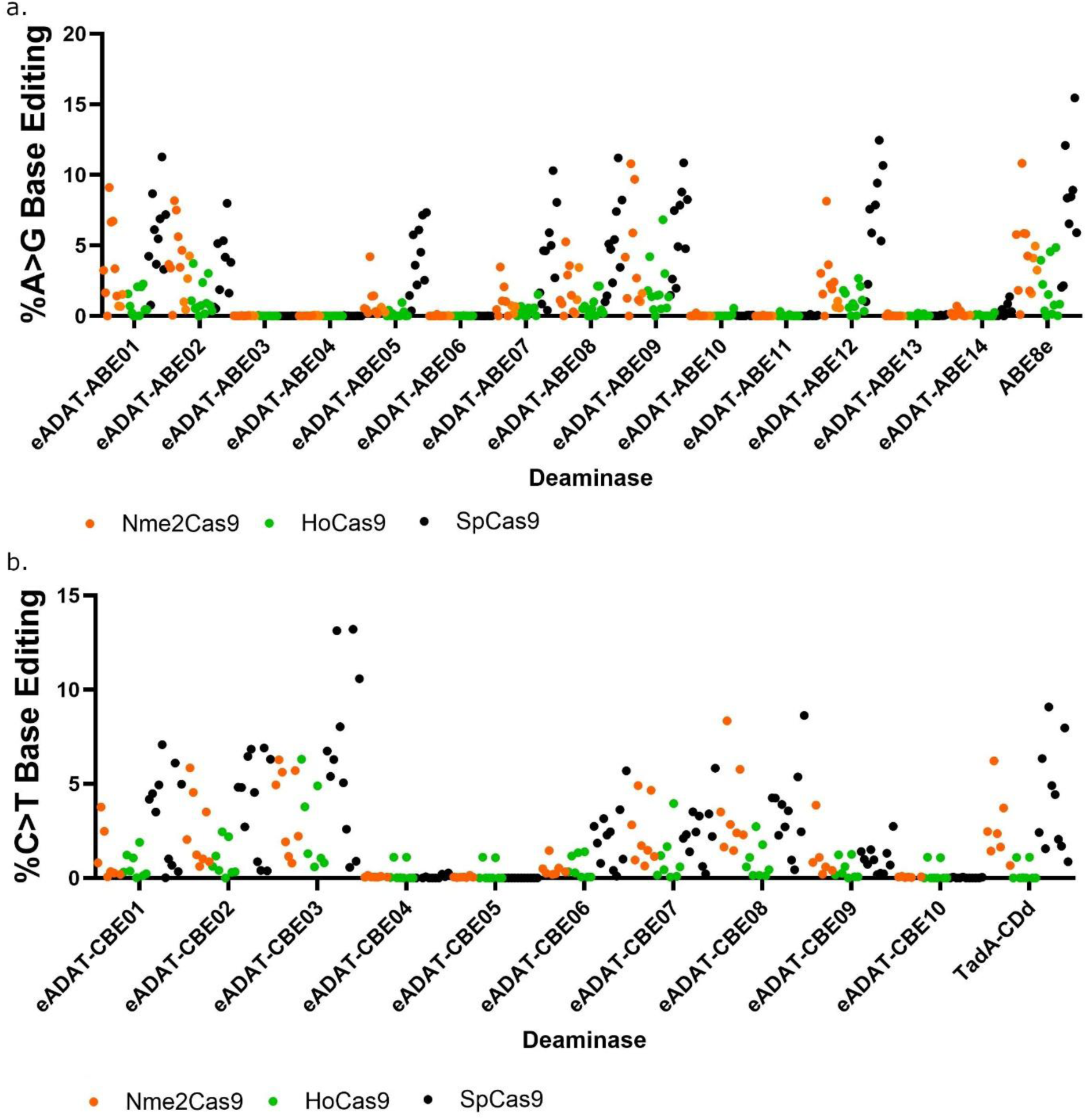
Eukaryotic activity of adenosine (a) or cytosine (b) deaminases identified from directed evolution across multiple targets and RGNs. Each point represents a single measurement of an independent target.

**Supplemental Figure 11.**
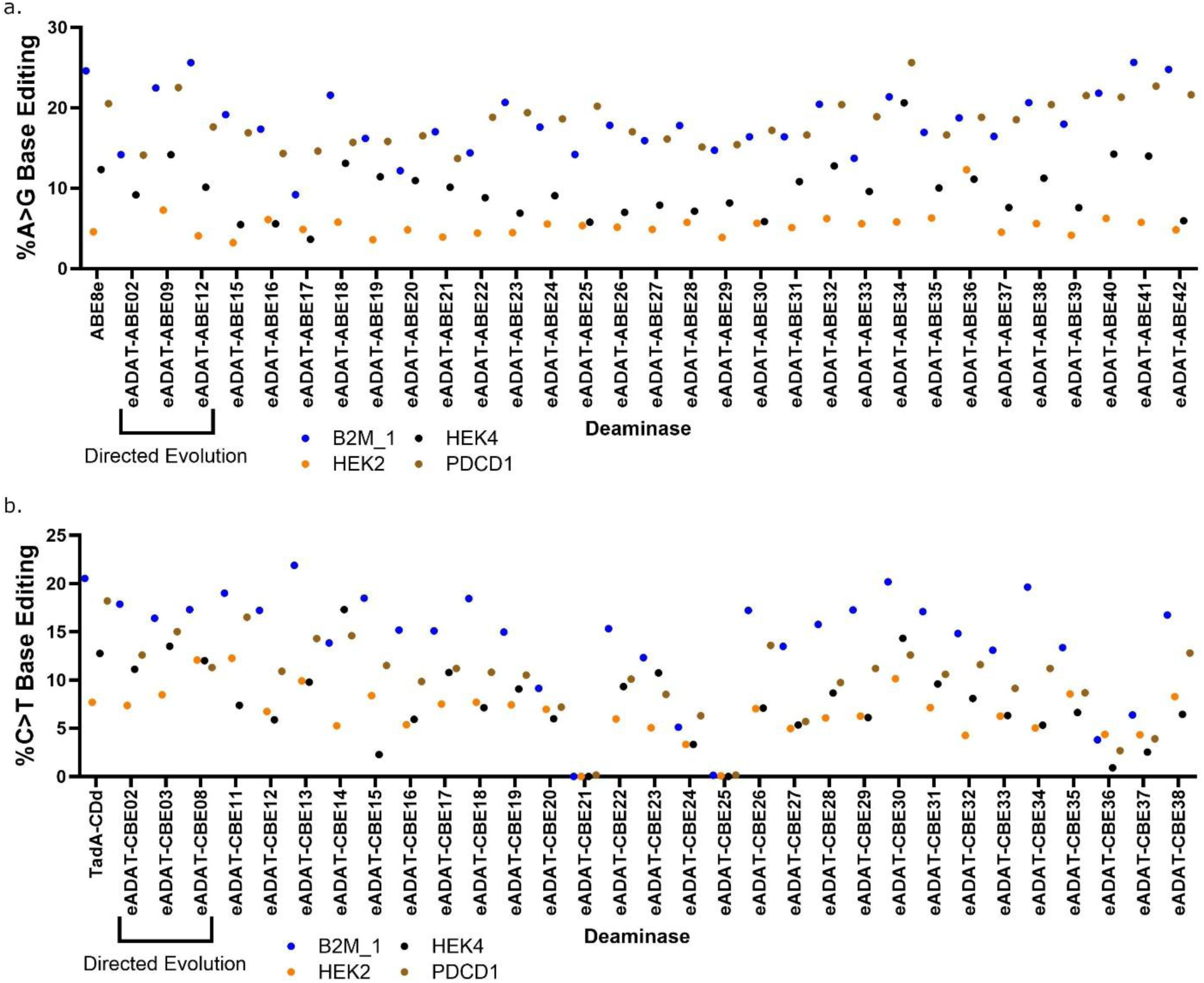
Eukaryotic activity of adenosine (a) or cytosine (b) deaminases identified from machine learning across four targets. Each point represents a single measurement of an independent target. ABEs are shown on top and CBEs are shown on bottom

**Supplemental Figure 12.**
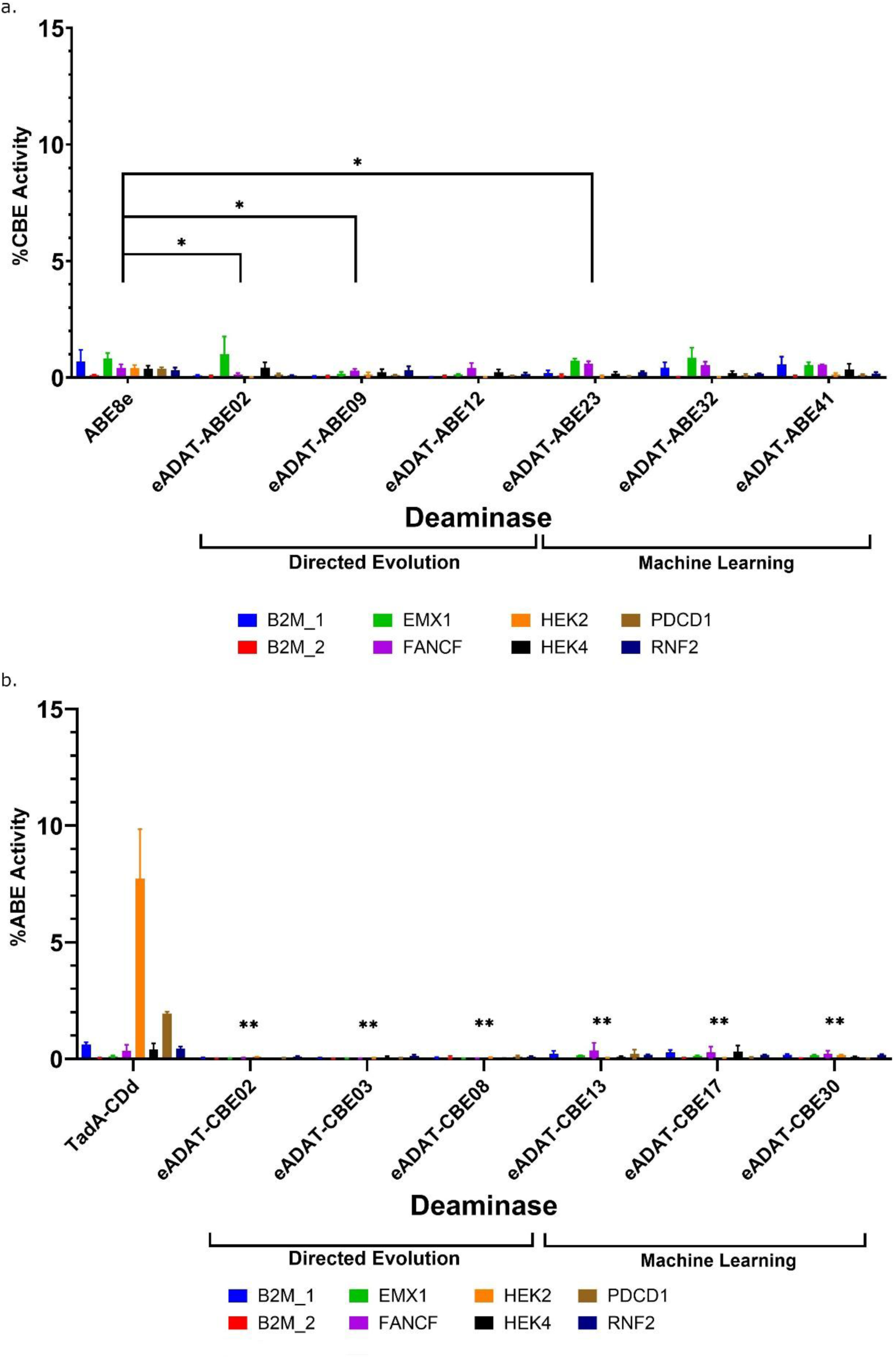
Off Base Target Eukaryotic Validation of Top Deaminases. Mean of the replicates is shown and Error bars represent the standard error of the mean. *=p<0.05, **=p<0.005, ***=p<0.0005, ****=p<0.00005. n=3 replicates for experimental conditions and n=5 replicates for controls. (a) Cytosine deamination with top adenosine deaminases across eight targets. Statistically significant differences from ABE8e are indicated, and all remaining comparisons to ABE8e showed no statistically significant difference. (b) Adenosine deamination with top cytosine deaminases across eight targets. Statistically significant differences from TadA-CDd are indicated.

**Supplemental Figure 13.**
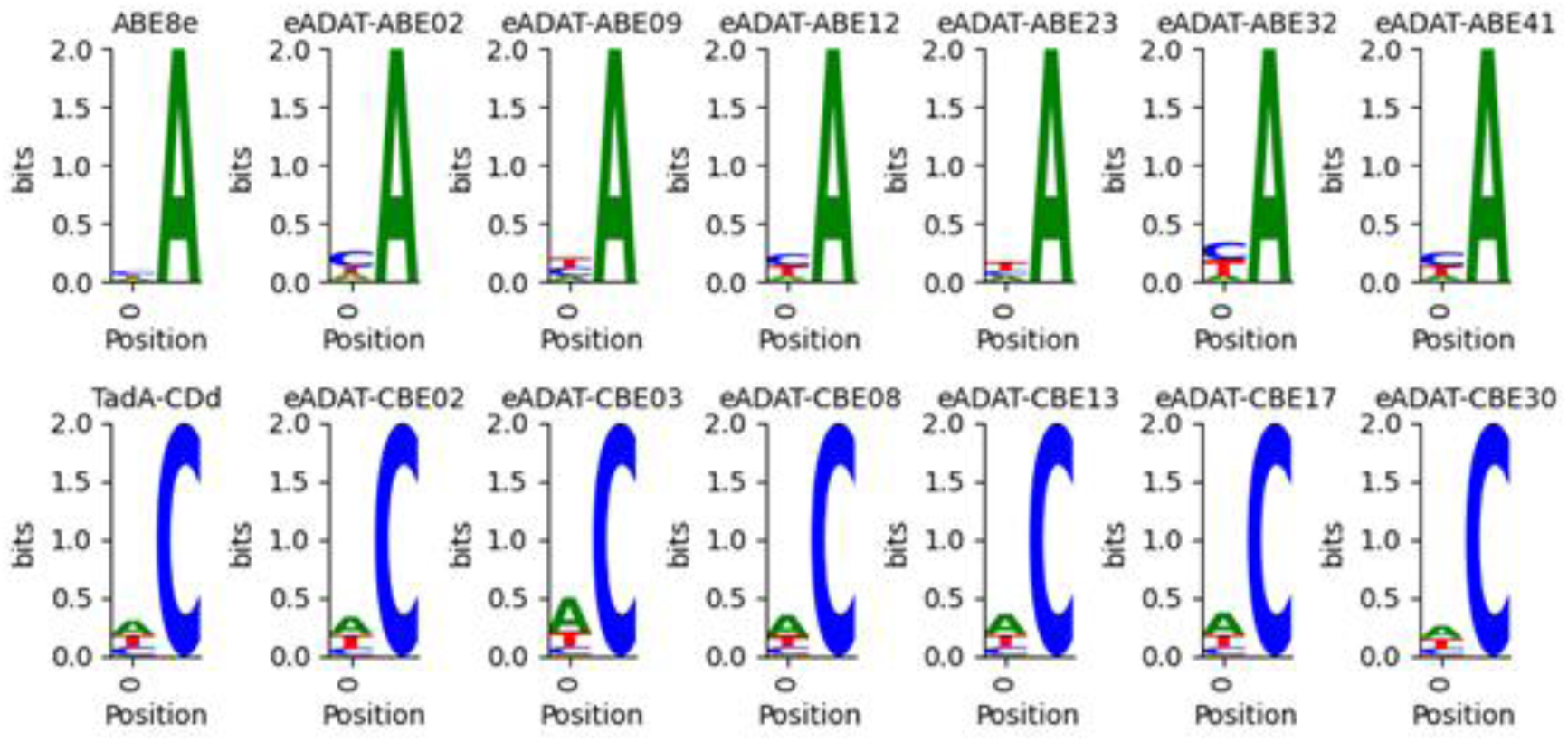
Dinucleotide context of edited adenosine (top) or cytosine (bottom) in DNA across eight different targets with n=3 replicates for experimental conditions and n=5 replicates for controls

**Supplemental Figure 14.**
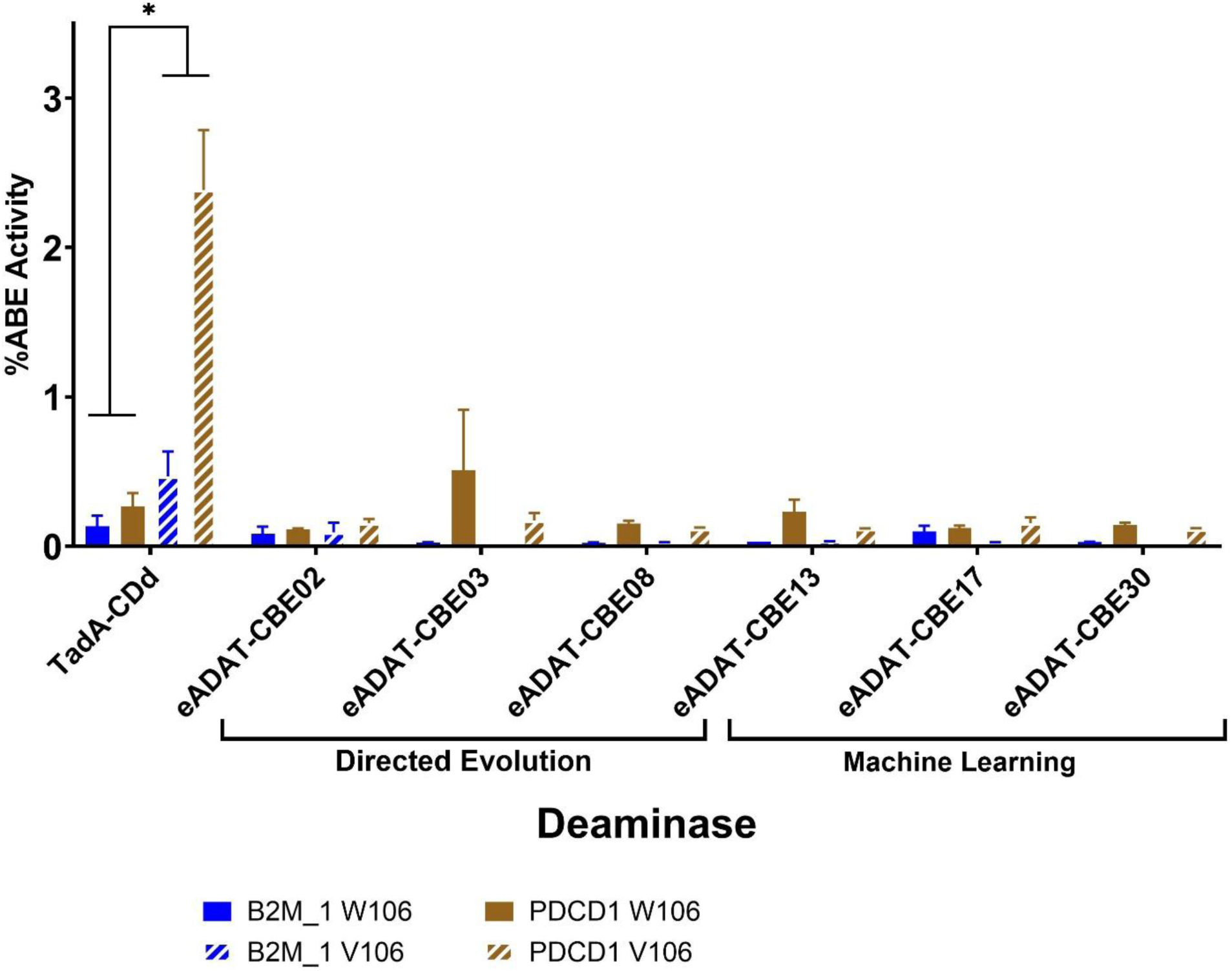
Adenosine off-base activity of CBE deaminases with valine or tryptophan at homologous residue of ecTadA 106. Bars represent the mean of n=3-4 replicates for all conditions. Error bars represent the standard error of the mean. Statistically significant differences between the Valine or Tryptophan residue in a construct are indicated. Remaining comparisons between mutants showed no statistically significant difference. *=p<0.05, **=p<0.005, ***=p<0.0005,****=p<0.00005

**Supplemental Figure 15.**
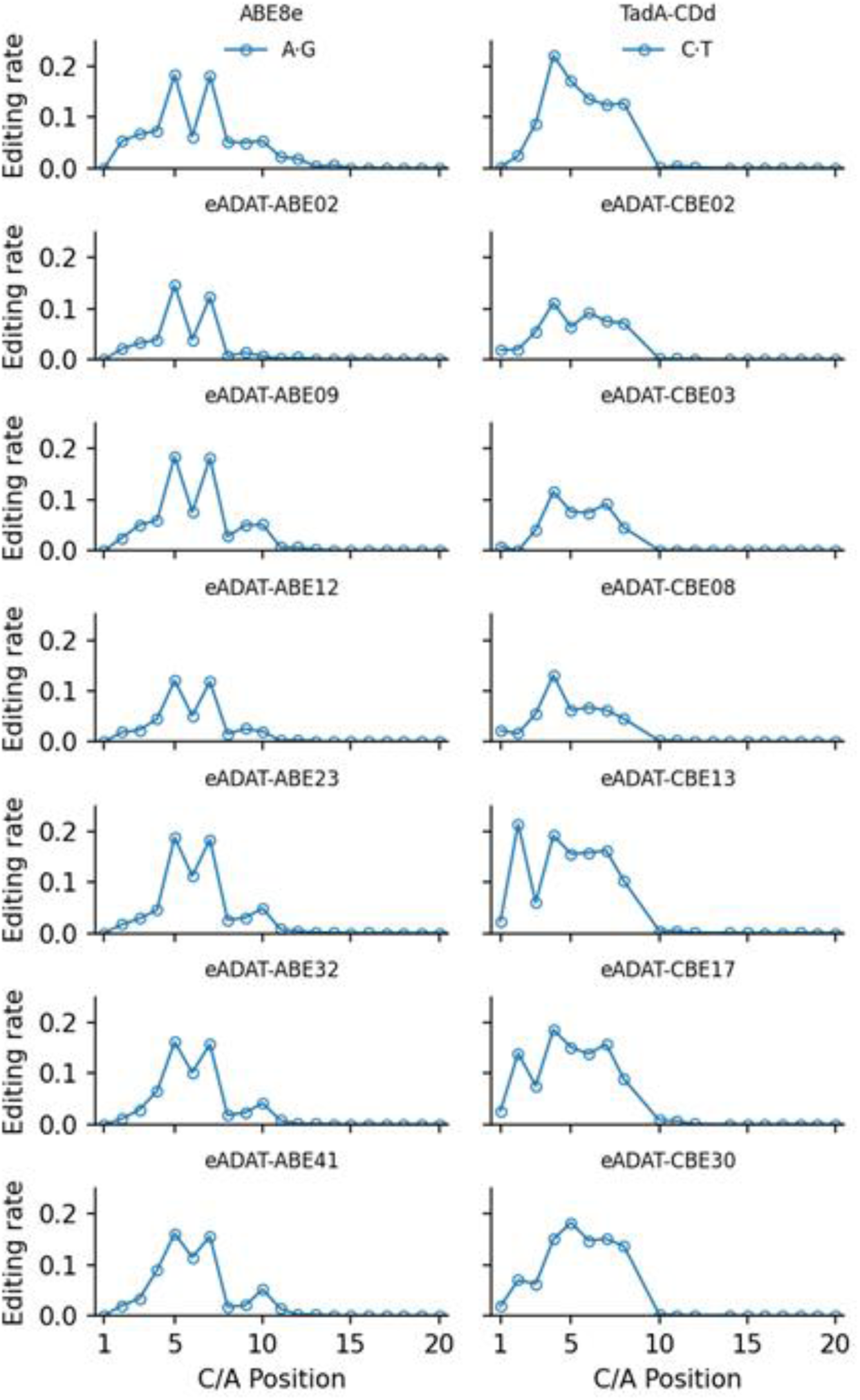
Editing window of ABE (left) or CBE (right) within the target sequence across eight different targets with n=3 replicates for experimental conditions and n=5 replicates for controls

**Supplemental Figure 16.**
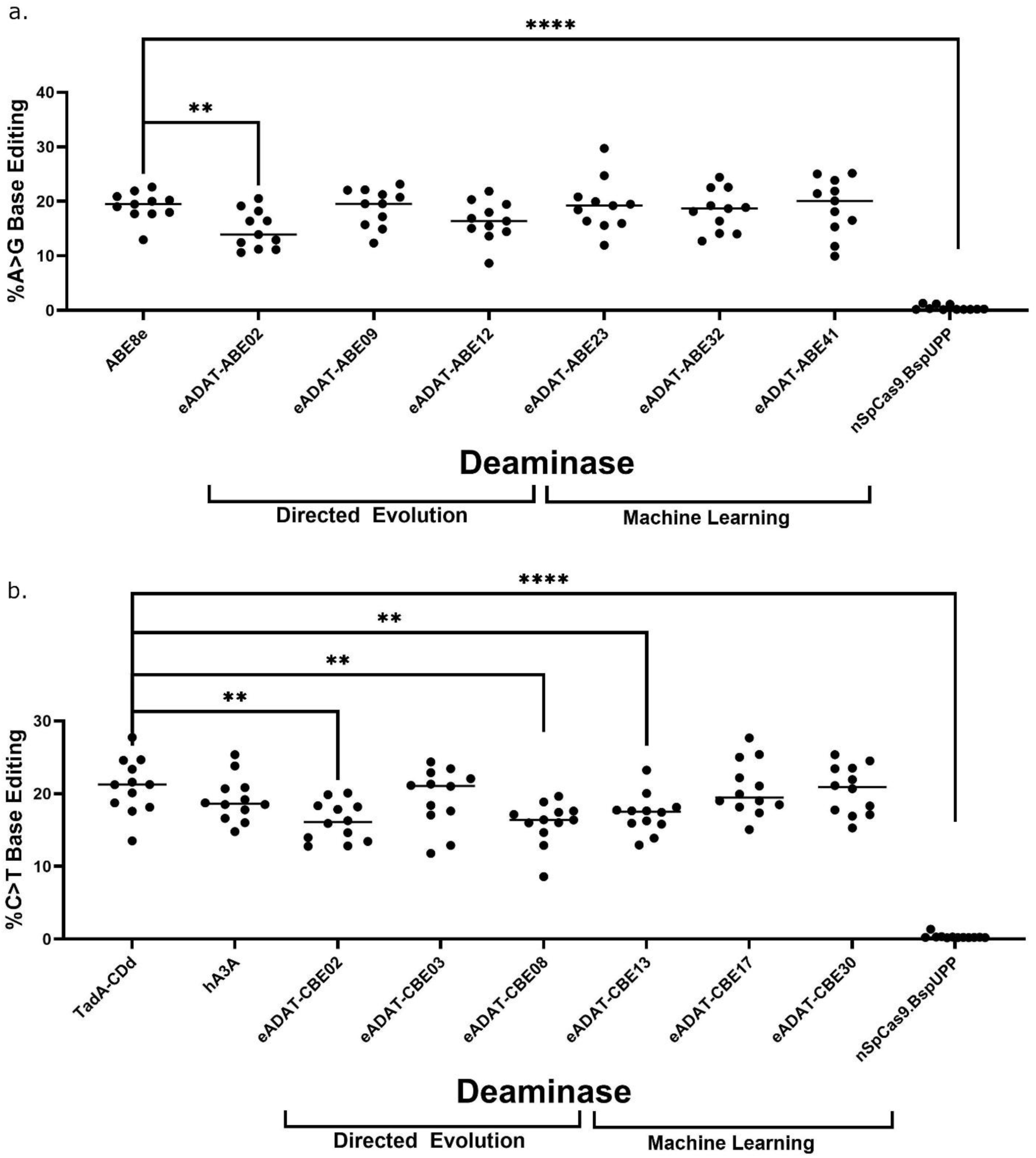
Eukaryotic on target activity at B2M_01 target of top adenosine (a) or cytosine (b) deaminases for off-target orthogonal R-loop assay. Bars represent the mean of n=11-12 replicates. Statistically significant differences from control deaminase are indicated.

**Supplemental Figure 17.**
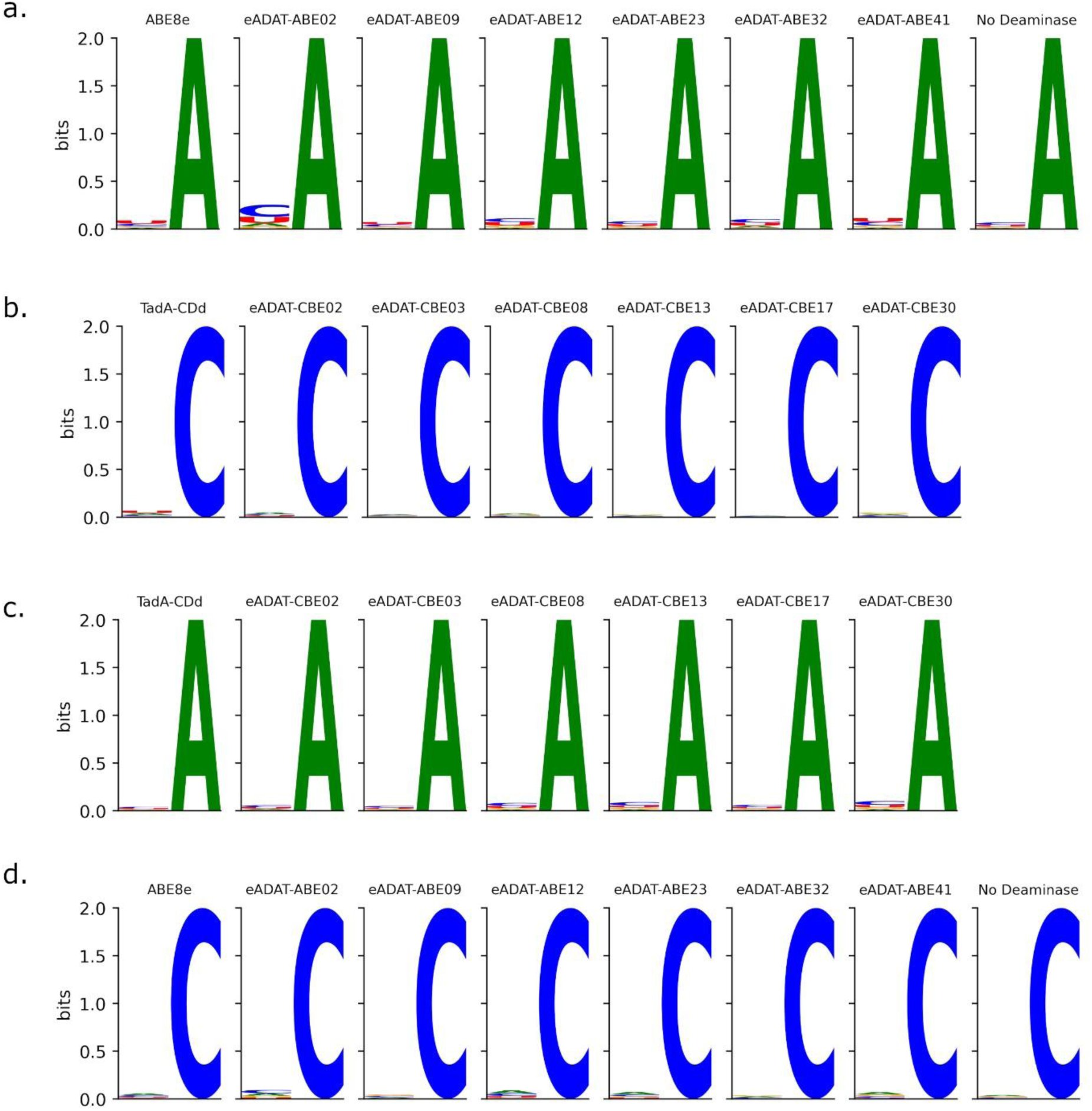
Dinucleotide context of deamination SNV in RNA. (a) Dinucleotide context of edited Adenosines by top ABEs. (b) Dinucleotide context of edited Cytosines by top CBEs. (c) Dinucleotide context of edited Adenosines by top CBEs. (d) Dinucleotide context of edited Cytosines by top ABEs.

**Supplemental Figure 18.**
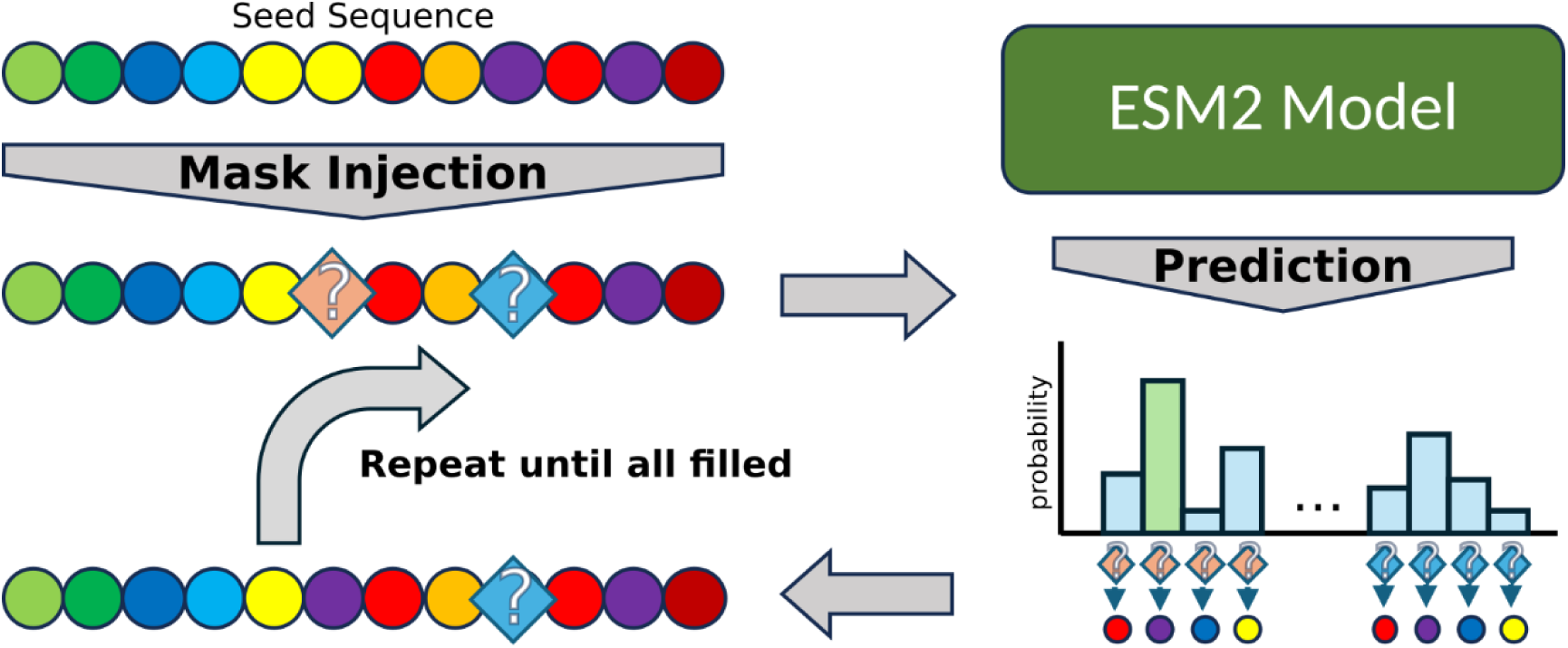
ESM mask filling sequence generation approach

## Supplementary Tables

We provide them as an xlsx file at bioRxiv.

## Supplementary Code

We provide this as a pdf file at bioRxiv.

## Notes

### Summary of Updates

The manuscript has been updated through corrections of inaccuracies, clarifications, and minor refinements of the experimental work and analyses.

