## Supplementary Code for "Machine Learning-Driven Optimization of Specific, Compact, and Efficient Base Editors via Single-Round Diversification"

### acbe_minimum_code

May 27, 2025

#### 0.1 train_multitask.py

[ ]:
```
from posixpath import abspath
from random import weibullvariate
from turtle import pos
import torch
import numpy as np
from tqdm import tqdm

from models import get_model
from torch.utils.data import DataLoader
from torch.nn.utils.rnn import pad_sequence

from torch import nn
from torch import optim

from utils.config import load_master_config
from utils.training import train_episode, validate, EarlyStoppingTracker,␣
↪_forward_result, reduce_masked_loss, _forward
from utils.seeding import fix_seeds
from data import get_dataset

import yaml, os

from torch.utils.tensorboard import SummaryWriter
from sklearn.metrics import roc_auc_score, average_precision_score

run_config_use = os.environ['USE_CONFIG']
n_folds = os.environ['HP_NFOLDS'] if 'HP_NFOLDS' in os.environ else 1
seed = int(os.environ['RNG_SEED'])

config = load_master_config(run_config_use, substitute_vars=True) training_conf␣
↪= config['training']
model_conf = config['model']

fix_seeds(int(training_conf['rng_seed']))

val_ds = get_dataset(**training_conf['val_ds'])
```

```python
train_ds = get_dataset(**training_conf['train_ds'])
test_ds = get_dataset(**training_conf['test_ds'])

def collate_fn_seq(data):
    xs = [d[0] for d in data]
    ys = [d[1] for d in data]
    return pad_sequence(xs, batch_first=True), torch.vstack(ys)
t_dataloader = DataLoader(
    train_ds,
    collate_fn=collate_fn_seq,
    shuffle=True,
    **training_conf['dls']
)
v_dataloader = DataLoader(
    val_ds,
    collate_fn=collate_fn_seq,
    shuffle=False,
    **training_conf['dls']
)
te_dataloader = DataLoader(
    test_ds,
    collate_fn=collate_fn_seq,
    shuffle=False,
    **training_conf['dls']
)

from torch.nn.functional import binary_cross_entropy_with_logits

class BCEMasked(nn.Module):
    def __init__(self, weights, reduction='mean', masked_value=-100):
        super(BCEMasked, self).__init__()

        self.weights = weights
        self.mv = masked_value
        self.reduction = reduction

    def forward(self, input, target):
        mask = target!=self.mv
        l = binary_cross_entropy_with_logits(input, target.to(dtype=torch.
 ↪float), pos_weight=self.weights, reduction='none')*mask.long()
        l = l.mean(0).sum()
        return l

model = get_model(**model_conf).cuda()

tb_sw = SummaryWriter(log_dir=training_conf['log_dir'], flush_secs=10) print(os.
 ↪path.join(training_conf['log_dir'], 'running_config.yaml'))
```

```
with open(os.path.join(training_conf['log_dir'], 'running_config.yaml'), 'wt')␣
↪as f:
    yaml.dump(config, f)

if os.environ.get('OPT_TYPE') is None or os.environ.get('OPT_TYPE') == "Adam":
    opt = optim.Adam(model.parameters(), **training_conf['opt'])
elif os.environ.get('OPT_TYPE') == "SGD":
    opt = optim.SGD(model.parameters(), **training_conf['opt'])
else:
    raise NotImplemented

weights = [(1-train_ds.y[:,i][train_ds.y[:,i]!=-100].mean(0))/train_ds.y[:
↪,i][train_ds.y[:,i]!=-100].mean(0) for i in range(train_ds.y.shape[1])]
weights = torch.tensor(weights).to(dtype=torch.float).cuda()
weights[weights<1.] = 1.

print(f'CE weights for classes:')
print(weights)
print(weights.shape)
loss = BCEMasked(weights=weights).cuda()

earlystopper = EarlyStoppingTracker(
    model=model,
    save_path=training_conf['log_dir'],
    tolerance=training_conf['train_tolerance_epochs'],
    more_better=True,
    improvement_threshold=1e-3,
)

@torch.no_grad()
def collect_preds(model , dl):
    ys = []
    yhats = []
    for x,y in dl:
        y_hat = _forward_result(x, model)
        ys.append(y)
        yhats.append(y_hat)
    return torch.cat(yhats, dim=0).cpu(), torch.cat(ys, dim=0).cpu()

def compute_taskwise_auc(yhats, ys, maskval = -100):
    yhats = yhats.numpy()
    ys = ys.numpy()
    AUC_mean = 0.
    AUCs = []
    for j in range(ys.shape[-1]):
        v_ys_subset = ys[:,j]
        v_yhats_subset = yhats[:,j]
```

```
        mask = v_ys_subset!=maskval
        AUCs.append(roc_auc_score(v_ys_subset[mask], v_yhats_subset[mask]).
 ↪item())
        AUC_mean += AUCs[-1]/ys.shape[-1]
    return AUCs, AUC_mean

def compute_taskwise_aupr(yhats, ys, maskval = -100):
    yhats = yhats.numpy()
    ys = ys.numpy()
    AUPR_mean = 0.
    dAUPR_mean = 0.
    dAUPRs = []
    AUPRs = []
    for j in range(ys.shape[-1]):
        v_ys_subset = ys[:,j]
        v_yhats_subset = yhats[:,j]
        mask = v_ys_subset!=maskval

                n_actives = (v_ys_subset[mask] == 1).sum()
        n_inactives = (v_ys_subset[mask] == 0).sum()
        n_total = n_actives + n_inactives

                random_clf_approx = n_actives / n_total
        auprc = average_precision_score(v_ys_subset[mask],␣
 ↪v_yhats_subset[mask]).item()

                AUPRs.append(auprc)
        dAUPRs.append(auprc - random_clf_approx)
        AUPR_mean += AUPRs[-1]/ys.shape[-1]
        dAUPR_mean += dAUPRs[-1]/ys.shape[-1]

    return AUPRs, dAUPRs, AUPR_mean, dAUPR_mean

if testing_mode:
        training_conf['max_epochs'] = 1

try:
    tb_total_steps = 0
    for i in range(training_conf['max_epochs']):
        val_loss, _ = validate(
            model,
            loss,             v_dataloader,
        )

        with torch.no_grad():
            val_loss = torch.stack(val_loss).mean()
            tb_sw.add_scalar('model/val_loss', val_loss, tb_total_steps)
```

```
        val_yhats, val_ys = collect_preds(model, v_dataloader)
        AUCs, AUC_mean = compute_taskwise_auc(val_yhats, val_ys)
        AUPRs, dAUPRs, AUPR_mean, dAUPR_mean = compute_taskwise_aupr(val_yhats,␣
↪val_ys)

                [tb_sw.add_scalar(f'model/AUC_task_{i}', aucval,␣
↪tb_total_steps) for i, aucval in enumerate(AUCs)]
        [tb_sw.add_scalar(f'model/AUPR_task_{i}', aucval, tb_total_steps) for␣
↪i, aucval in enumerate(AUPRs)]
        [tb_sw.add_scalar(f'model/dAUPR_task_{i}', aucval, tb_total_steps) for␣
↪i, aucval in enumerate(dAUPRs)]
        tb_sw.add_scalar(f'model/mean_AUPR', AUPR_mean, tb_total_steps)
        tb_sw.add_scalar(f'model/mean_AUC', AUC_mean, tb_total_steps)
        tb_sw.add_scalar(f'model/mean_dAUPR', dAUPR_mean, tb_total_steps)

                [print(f'{i} - {AUCs[i]} - {AUPRs[i]} - {dAUPRs[i]}') for i in␣
↪range(len(AUCs))]

                if earlystopper.update(dAUPR_mean):
            print(f'Earlystopping: max duration without improvement exceeded.')
            break

        print(f"Epoch {i}")
        train_losses, _ = train_episode(
            model,
            opt,
            loss,
            t_dataloader,
        )
        train_losses_epoch = torch.stack(train_losses).mean()
        print(f'Train loss: {train_losses_epoch}')
                        for l in train_losses:
            tb_sw.add_scalar('model/train_loss', l, tb_total_steps)
            tb_total_steps += 1
                tb_sw.add_scalar('model/train_loss', train_losses_epoch,␣
↪tb_total_steps)

except KeyboardInterrupt as e:
    print('KBINT')
finally:
    earlystopper.finalize()
    model = earlystopper.best_model().cuda()
    print('Dumping validation predictions for the best model...')

    val_yhats, val_ys = collect_preds(model, v_dataloader)
```

```
    AUCs, AUC_mean = compute_taskwise_auc(val_yhats, val_ys)
    AUPRs, dAUPRs, AUPR_mean, dAUPR_mean = compute_taskwise_aupr(val_yhats,␣
↪val_ys)
        reloss = BCEMasked(weights=torch.ones_like(val_yhats)[0, :]).cuda()
    with torch.no_grad():
        val_reloss = reloss(val_yhats, val_ys).cpu()

    print(f'Mean validation AUC of the final model: {AUC_mean}, unweighed loss␣
↪of {val_reloss.item()}')
    resdic = {
        'val_AUC': AUCs,
        'val_mean_AUC': AUC_mean,          'val_AUPR': AUPRs,
        'val_mean_AUPR': AUPR_mean,          'val_dAUPR': [a.item() for a in␣
↪dAUPRs],
        'val_mean_dAUPR': float(dAUPR_mean),          'val_unweighed_loss':␣
↪val_reloss.item(),
    }

        test_yhats, test_ys = collect_preds(model, te_dataloader)
    AUCs, AUC_mean = compute_taskwise_auc(test_yhats, test_ys)
    AUPRs, dAUPRs, AUPR_mean, dAUPR_mean = compute_taskwise_aupr(test_yhats,␣
↪test_ys)
    with torch.no_grad():
        te_reloss = reloss(test_yhats, test_ys).cpu()

    print(f'Mean test AUC of the final model: {AUC_mean}, unweighed loss of␣
↪{te_reloss.item()}')
    resdic['test_AUC'] = AUCs
    resdic['test_mean_AUC'] = AUC_mean     resdic['test_AUPR'] = AUPRs
    resdic['test_mean_AUPR'] = AUPR_mean     resdic['test_dAUPR'] = [a.item()␣
↪for a in dAUPRs]
    resdic['test_mean_dAUPR'] = dAUPR_mean.item()
    resdic['test_unweighed_loss'] = te_reloss.item()

    with open(os.path.join(training_conf['log_dir'], 'best_model_performance.
↪yaml'), 'wt') as f:
        yaml.dump(resdic, f)

    print('done.')
```

##### 0.2 cnn_activity_regressor.py

[ ]:
```
import torch
from torch import nn

class CNNActivityRegressor(nn.Module):
```

```python
    def __init__(
        self,
        layers: list,
        linear_params: dict,
        pool_dim: int = 12,
        act = nn.ELU(),
        aa_number: int = 22,
        pooling='mean',
        p_dropout=0.2,
        posenc_enabled='none',
        pooling_kwargs = {},
        ):

        super(CNNActivityRegressor, self).__init__()

        self.convs = nn.ModuleList([nn.Conv1d(**p) for p in layers])
        self.output = nn.Linear(**linear_params)
        self.act = act
        self.dropout = nn.Dropout(p=p_dropout)

        if pooling=='mean':
            self.pooling = nn.AdaptiveAvgPool1d(**pooling_kwargs)
        elif pooling=='max':
            self.pooling = nn.AdaptiveMaxPool1d(**pooling_kwargs)
        elif pooling=='lstm':
            self.pooling = nn.LSTM(**pooling_kwargs)
        elif pooling=='dotprod':
            raise NotImplemented
        else:
            raise ValueError

        self.aa_number = aa_number
        if posenc_enabled=='none':
            pass
        elif posenc_enabled=='learned':
            self.posenc = nn.Embedding(num_embeddings=1024, embedding_dim=aa_number, max_norm=1.)
        else:
            raise ValueError

    def _one_hot_encode(self, x):
        x = x.to(dtype=torch.long)
        x = nn.functional.one_hot(x, num_classes=self.aa_number+1).to(device=next(self.parameters()).device, dtype=next(self.parameters()).dtype)[:,:,1:]
        x = torch.permute(x, (0, 2, 1))
        return x
```

```python
    def forward(self, x):
        if x.ndim<3:
            x = self._one_hot_encode(x)

        assert x.ndim==3, f"Dimension of the output is {x.ndim}, gotta be 3."

        if hasattr(self, 'posenc'):
            x += self.posenc(torch.arange(0, x.shape[2]).to(device=x.device)).
↪unsqueeze(0).expand(x.shape[0], -1, -1).permute(0, 2, 1)

        for conv in self.convs:
            x = conv(x)
            x = self.act(x)

        x = self.pooling(x) if not isinstance(self.pooling, nn.LSTM) else self.
↪pooling(x)[0]

        x = self.dropout(x)
        x = torch.flatten(x, start_dim=1,end_dim=2)

        x = self.output(x)
        return x
```

##### 0.3 esm_generation.py

[ ]:
```python
"""
An ESM model is used for sequence generation for the CBE-ADATS.

General idea:
 * Start with a known sequence
 * Mask some amino acids
 * Let the ESM model replace them and end up with a new sequence

The generation process includes following steps:
 (1) Seed sampling
 (2) Mask sampling
 (3) Iterative mask filling

In a pre-processing step we extracted the top k (k=32; design choice) sequences
for every rgn which resulted in a list of 95 unique sequences. For each mini-
batch with sample a sequence from this list as a starting point for the
generation process. We call this starting point seed.

For each mini-batch we uniformly sample 1 to 5 mask postitions. (Range 1-5:
design choice).
```

```python
Assuming the mask sampling step returned i mask positions, we run i iterations:
 * We feed the masked sequences into the ESM-model and return the logits for the
   masked position. This gives a distribution across possible amino acids for
   each masked position.
 * We create a joint distribution by merging the distributions for the different
   positions which results in a sampling distrubtions for tuples (pos, aa).
 * From the joint dristibution we sample a position and replace the mask with
   the sampled amino acid.
 * We repeat this until all masks are filled
"""

INPUT_PATH_TRAIN_SEQS = ("/home/asceusr/blob-storage/jku/data_oct23/cd-adat/"
                         "results/raw_counts.txt")
INPUT_PATH_SEEDS = ("/home/asceusr/blob-storage/jku/screening_11_23/"
                    "top_k_seeds_for_generation/seeds_cbe.npy")
OUTPUT_PATH = ("/home/asceusr/blob-storage/jku/data_oct23/generated_seqs/"
               "cd-adat/esm/with_deletions/v_01/generated_seqs.pkl")
BATCH_SIZE = 20

import esm
import torch
import numpy as np
import pandas as pd
import copy
import pickle


class SequenceGenerator:
    """
    This class generates sequences by
    (1) loading the esm model,
    (2) seed sampling
    (3) injecting mutations into the consensus sequence,
    (4) sampling from the mask distribution to fill the masked positions,
    (5) repeating (4) until all masked positions are filled.
    """

    def __init__(self, already_generated_seqs=None):
        """
        already_generated_sequences: list
        """

        self.model, self.alphabet = esm.pretrained.esm2_t33_650M_UR50D()
        self.batch_converter = self.alphabet.get_batch_converter()
        self.model.cuda()
```

```python
        self.id_to_tok_dict = {i: tok for i, tok in
                               enumerate(self.alphabet.all_toks)}
        self.tok_to_id_dict = {tok: i for i, tok in
                               enumerate(self.alphabet.all_toks)}
        self.standard_tok_ids = [self.tok_to_id_dict[tok] for tok in
                                 self.alphabet.standard_toks]

        self.id_mask = self.tok_to_id_dict["<mask>"]

        data = pd.read_csv(INPUT_PATH_TRAIN_SEQS, sep='\t')
        train_seqs = list(data.iloc[:,0].values)
        if already_generated_seqs != None:
            train_seqs = train_seqs + already_generated_seqs
        self.train_seqs = tuple(train_seqs)

        self.seeds = np.load(INPUT_PATH_SEEDS)
        self.nbr_seeds = self.seeds.shape[0]

        self.sequence_buffer = set()

        self.min_nbr_mutations = 1
        self.max_nbr_mutations = 5

        self.batch_size = BATCH_SIZE

    def generate_new_sequence(self):
        """
        This function generates new sequences by
        (1) Seed sampling
        (2) Defining the nbr of mutations
        (3) Injecting mutations
        (4) Sampling from the mask distribution to fill the masks
        (5) Repeating (4) until all masks are filled
        """
        def sample_mutation_positions(seed_length:int, nbr_mutations:int,
                                      batch_size=BATCH_SIZE) -> list:
            """
            This function samples (without replacement) positions where
            mutations should be injected
            """

            sampled_positions = list()

            for _ in range(batch_size):
                pos = torch.randperm(seed_length)[:nbr_mutations]
                pos = pos.sort()[0]
                sampled_positions.append(pos)
```

```
            return sampled_positions

        def inject_mutations_by_masking(sequence:str,
                                        positions:list,
                                        mask_token:str='<mask>',
                                        batch_size=BATCH_SIZE) -> list:
            """
            This function injects a mutation into a sequence by masking the␣
↪positions
            """

            masked_sequences = list()

            for i in range(batch_size):

                masked_sequence = copy.deepcopy(sequence)

                pos_bias = 0
                for pos in positions[i]:
                    real_pos = pos + pos_bias
                    masked_sequence = (masked_sequence[:real_pos] +
                                       mask_token +
                                       masked_sequence[real_pos+1:])
                    pos_bias += len(mask_token) - 1

                masked_sequences.append(masked_sequence)

            return masked_sequences

        def fill_mask_position(esm_logits:torch.Tensor,
                               mask_positions:torch.Tensor,
                               batch_size=BATCH_SIZE):

            def one_seq_sample_masked_position_and_filling_from_sampling_distr(
                sampling_distr:torch.Tensor,):
                """
                This function samples a masked position and fills it with a AA␣
↪sampled
                from the given distributions
                """

                sampling_distr_dim0, sampling_distr_dim1 = sampling_distr.shape

                candidates_index = np.zeros(sampling_distr_dim0 *␣
↪sampling_distr_dim1 )
                candidates = list()
```

```
                c_index = 0
                for p in list(range(sampling_distr_dim0)):
                    for alphabet_id in range(sampling_distr_dim1):
                        candidates_index[c_index] = c_index
                        c_index += 1
                        candidates.append((p, alphabet_id))

                merged_distr = sampling_distr.reshape(1,
                                                      sampling_distr_dim0 *
                                                      sampling_distr_dim1)
                merged_distr = torch.nn.Softmax(dim=1)(merged_distr).numpy().
↪flatten()
                repeat_drawing = True                 while repeat_drawing:
                    drawn_c_index = np.random.choice(candidates_index, size=1,␣
↪p=merged_distr)
                    drawn_candidate = candidates[int(drawn_c_index)]

                    drawn_masked_id, drawn_alphabet_id = drawn_candidate
                    if drawn_alphabet_id != 32:                             ␣
↪repeat_drawing = False

                return drawn_masked_id, drawn_alphabet_id

            drawn_masked_ids = list()
            drawn_alphabet_ids = list()
            for i in range(batch_size):
                sampling_distr = torch.squeeze(esm_logits[i, mask_positions[i],
                                                           :])
                if len(sampling_distr.shape) == 1:
                    sampling_distr = sampling_distr.reshape(1, -1)

                assert len(sampling_distr.shape) == 2
                sampling_distr = torch.nn.Softmax(dim=1)(sampling_distr)
                (drawn_masked_id, drawn_alphabet_id
                 ) =␣
↪one_seq_sample_masked_position_and_filling_from_sampling_distr(
                    sampling_distr=sampling_distr)

                drawn_masked_ids.append(drawn_masked_id)
                drawn_alphabet_ids.append(drawn_alphabet_id)

            return drawn_masked_ids, drawn_alphabet_ids

        def create_sequence_representations_from_tokens(token_ids:torch.Tensor,
                                                        alphabet:esm.Alphabet,
                                                        standard_tok_ids:list,
                                                        batch_size=BATCH_SIZE):
```

```
        """
        This function creates sequence representations from tokens
        """

        def process_one_sequence(token_ids,
                                 alphabet,
                                 standard_tok_ids):
            token_ids = torch.squeeze(token_ids)

            sequence = ""
            for token_id in token_ids:
                if token_id.item() in standard_tok_ids:
                    sequence += alphabet.get_tok(token_id.item())

            return sequence

        sequences = list()

        for i in range(batch_size):
            tokens = token_ids[i,:]
            sequence = process_one_sequence(tokens,
                                            alphabet,
                                            standard_tok_ids)
            sequences.append(sequence)

        return sequences

    sampled_seed_id = np.random.randint(0, self.nbr_seeds)
    seed = self.seeds[sampled_seed_id]

    nbr_deletions = np.random.randint(1,4)
    for _ in range(nbr_deletions):
        i = np.random.randint(1,len(seed)-1)
        seed = seed[:i] + seed[i+1 :]

    seed_length = len(seed)

    nbr_mutations = np.random.randint(self.min_nbr_mutations,
                                      self.max_nbr_mutations)

            sampled_positions = sample_mutation_positions(seed_length,
                                                          nbr_mutations)

    masked_sequences = inject_mutations_by_masking(seed, sampled_positions)

            data = list()
    for i, masked_sequence in enumerate(masked_sequences):
```

```
            data.append((f"p_{i}", masked_sequence))
        _, _, batch_tokens = self.batch_converter(data)

                while self.id_mask in batch_tokens:

                        mask_positions = torch.squeeze(batch_tokens == self.
↪id_mask)
            mask_position_ids = list()
            for i in range(self.batch_size):
                ids = torch.where(batch_tokens[[i], :] == self.id_mask)[1]
                mask_position_ids.append(ids)

                                    with torch.no_grad():
                results = self.model(batch_tokens.cuda(),
                                     repr_layers=[33],
                                     return_contacts=True)

            logits = results['logits'].detach().cpu()

                        ( drawn_masked_ids, drawn_alphabet_ids
            ) = fill_mask_position(logits, mask_positions)

                        for i in range(self.batch_size):
                batch_tokens[i,
                             mask_position_ids[i][drawn_masked_ids[i]]
                            ] = drawn_alphabet_ids[i]

                sequences =␣
↪create_sequence_representations_from_tokens(batch_tokens,
                                                            self.alphabet,
                                                            self.
↪standard_tok_ids)

                for sequence in sequences:
            if (sequence not in self.train_seqs and
                sequence not in self.sequence_buffer):
                self.sequence_buffer.add(sequence)

    def run_generation(self,
                       saving_dir:str,
                       saving_interval:int=3000,
                       max_nbr_sequences:int=999999999999999999):
        """
        This function generates sequences in a infite loop
        """

        continue_generation = True
```

```
        mini_batch_i = 0
        while continue_generation:
            self.generate_new_sequence()

            if mini_batch_i % saving_interval == 0:
                print(f'{len(self.sequence_buffer)} seqs generated ...')
                with open(saving_dir, 'wb') as f:
                    pickle.dump(self.sequence_buffer, f)

            if len(self.sequence_buffer) >= max_nbr_sequences:
                continue_generation = False

            mini_batch_i = mini_batch_i + 1

                print(f'End. {len(self.sequence_buffer)} seqs generated!')
        with open(saving_dir, 'wb') as f:
            pickle.dump(self.sequence_buffer, f)

if __name__ == "__main__":

    sequence_generator = SequenceGenerator()
    sequence_generator.run_generation(saving_dir=OUTPUT_PATH,
                                      saving_interval=1000,
                                      max_nbr_sequences=5000000)
```
